# Maternal high-fat, high-sucrose diet-induced excess adiposity is linked to placental hypoxia and disruption of fetoplacental immune homeostasis in late gestation

**DOI:** 10.1101/2024.09.30.615691

**Authors:** Christian J. Bellissimo, Tatiane A. Ribeiro, Erica Yeo, Patrycja A. Jazwiec, Howard Luo, Jaskiran Bains, Deborah M. Sloboda

## Abstract

Maternal excess adiposity (i.e., overweight and obesity) at conception is linked to numerous signs of malperfusion and inflammatory injury in the placenta. Previous reports have suggested that obesity-associated placental malperfusion may trigger a state of fetoplacental hypoxia, contributing to adverse health outcomes within and beyond the perinatal period. However, many previous studies have relied on indirect measures of tissue oxygen saturation, including readouts influenced by external inflammatory stressors. Direct comparisons of tissue oxygen saturation at the uteroplacental interface in pregnancies complicated by excess adiposity are lacking. Here, we used a mouse model of chronic preconception high-fat, high-sucrose (HFHS) diet feeding to model the impacts of an obesogenic milieu on placental oxygenation near term gestation (E17.5). We found that both placental junctional and labyrinth zone tissues were relatively hypoxic in HFHS pregnancies compared to chow-fed controls (CON). However, this was not associated with enhanced HIF-1α expression in labyrinth tissues. Similarly, placentas from CON and HFHS dams did not exhibit gross differences in morphology or vessel density and pericyte coverage. However, HFHS placentas have a greater burden of histopathological lesions, including tissue calcification and fibrinoid deposition within the labyrinth zone. Calcified placental tissue coincided with the destruction of vasculosyncytial membranes and macrophage-dense foci, alongside altered expression of immunomodulatory and chemotactic cytokines within the labyrinth zone proteome, which differed in magnitude with fetal sex. While fetal growth was not markedly affected, fetuses from HFHS pregnancies exhibited higher levels of circulating IL-6, prolactin, CXCL1, and CCL2. Collectively, these data confirm that diet-induced maternal excess adiposity leads to a relative state of placental hypoxia, even in the absence of marked growth restriction or fetal demise. While this hypoxic state is not linked to gross morphological abnormalities, it is associated with a greater histopathological burden indicative of local malperfusion and inflammation, and an altered fetal inflammatory and endocrine milieu in late gestation. These findings provide new insight into mechanisms by which an obesogenic environment during pregnancy compromises placental function and contributes to the long-term programming of chronic disease susceptibility.

## INTRODUCTION

Recent estimates report that nearly half of pregnant Canadians are overweight or obese at the time of conception (1, 2). Preconception excess adiposity is linked to extremes in offspring birth weight, which may arise secondary to other obstetrical complications that overweight or obese individuals are more predisposed to (3–5). This includes an elevated risk of gestational diabetes mellitus (GDM), preterm labour, or preeclampsia (6, 7). In turn, offspring exposed to this obesogenic environment *in utero* exhibit poorer cardiometabolic and neuropsychiatric outcomes throughout the life course (8–13). While the causal role of intrauterine adversity in the programming of chronic disease risk is now widely accepted, our understanding of mechanisms underlying adverse obstetric and postnatal outcomes in pregnancies impacted by maternal excess adiposity remains incomplete.

Accumulating evidence supports the notion that excess adiposity—particularly obesity—is associated with intrauterine hypoxia (14–21), adding to the growing constellation of endocrine (8), inflammatory (22), and cellular redox mechanisms (23) that prime for poor pre- and postnatal outcomes. Hypoxia *in utero* results from an imbalance between the aerobic demands of fetoplacental metabolism and the availability of oxygen supplied by maternal blood (24). It has been suggested that fetal hypoxia arises in a ‘post-placental’ manner in the context of maternal obesity, driven by enhanced oxygen demands because of high macronutrient delivery to the fetus, which stimulates growth factor secretion (e.g., insulin and insulin-like growth factors) and elevates fetal metabolic rate (8, 24, 25). Ectopic lipid deposition in fetal tissues in obese pregnancies may also contribute to a state of oxidative stress and inflammation, which may also increase fetal oxygen consumption (26–29).

However, recent evidence suggests that obesity can also compromise the supply of oxygen to the placenta by way of impaired maternal cardiovascular adaptations to pregnancy (6, 30–34), reduced uteroplacental blood flow (34, 35), or disruptions in the remodelling of decidual spiral arteries supplying placental tissue (36–38). Moreover, obesity is associated with structural placental changes that can impair the capacity for placental oxygen diffusion, including increases in interhaemal barrier thickness (29, 39) and inflammatory injury to villous tissue (18, 40, 41). This can independently reduce placental oxygen extraction from maternal blood (42). Indeed, placental histologic lesions of both maternal vascular malperfusion and chronic inflammation have been more commonly observed in human pregnancies complicated by obesity (18, 40, 43, 44). Collectively, reduced oxygen may induce angiogenic, endocrine, and/or inflammatory responses that modify fetal and placental development to maintain growth and survival at the expense of later postnatal health (45).

Changes to placental vascular development may be one consequence, with implications for both immediate and long-term offspring health. Placental vasculature is the principal conduit for nutrient and gas exchange and can adapt to changes in the supply of and demand for oxygen through both structural and molecular modifications (46–48). This adaptability is important for ensuring placental capacity for oxygen extraction is matched to the demands of fetal growth. However, changes in placental vascular development in response to hypoxic challenges may have collateral impacts on postnatal health. As a direct extension of the fetal cardiovascular system, changes in placental vascularization are thought to impact the development of the fetal heart (49–51). Indeed, several epidemiologic studies have described relationships between placental efficiency and morphometry and offspring risk of adverse cardiovascular outcomes (52–58). Altered offspring cardiac structure has also been described in rodent models of maternal diet-induced obesity (59–61).

Previous work in rodent models of preconception diet-induced obesity has reported increased placental vessel density and decreased microvascular stability, consistent with an angiogenic response to hypoxic stress (15–17, 19). Equally, others have reported reduced placental vascularization in similar models (62–67), and fetal growth outcomes associated with these phenotypes are just as variable (68). Interestingly, surrogate molecular markers of hypoxia within the placenta are elevated in the context of both increased and decreased vascular density with diet-induced metabolic impairments (14–17, 19, 67, 69). The exact cause(s) of these discrepant changes to placental vascularity in the face of apparent hypoxia are not fully understood.

Due to the common hemochorial mode of placentation shared with humans (70), rodent models of preconception obesogenic diet feeding are a common preclinical tool for studying fetoplacental adaptations to maternal excess adiposity and metabolic dysfunction (71). Unlike humans, however, the placentas of mice and rats are segregated into labyrinth and junctional zones, with the latter being perfused by oxygen-poor venous blood. This results in differential oxygen tension between these tissue layers and a greater susceptibility of junctional zone tissues to hypoxia (46, 72, 73). This fact has largely been unaccounted for in previous studies that rely on bulk tissue protein expression of Hypoxia-inducible factors (HIFs) as a readout of placental oxygenation. This has limited the ability to delineate the location of the hypoxia within the placenta, its relationship to consequences for the placental vascular interface, and determine its translational nature to human pregnancies.

Historical use of HIF-1α expression as a readout of tissue oxygenation is based on its canonical role as an oxygen-sensitive transcription factor that accumulates under hypoxic conditions (47). However, recent evidence has demonstrated that the stabilization and activity of HIFs are regulated by several other factors, including inflammatory cytokines and oxidative stress (47, 74). Inflammation and oxidative stress are both factors reported to be elevated in placentas of obese pregnancies (15–17, 26, 28, 39, 75). Excessive reactive oxygen species generated from increased mitochondrial beta-oxidation may inhibit HIF-degrading prolyl hydroxylase enzymes (76, 77), while inflammatory cytokines such as TNF may stimulate HIF transcription through induction of nuclear factor kappa B (NFκB) signalling (78, 79). Thus, total placental expression of HIF-1α or its transcriptional targets may not be a reliable measure of placental oxygenation where other oxidative and proinflammatory disruptions are present.

Understanding the relationship between fetoplacental oxygen supply and consequences for placental vascular adaptations in metabolically complicated pregnancies is key to developing strategic interventions to mitigate adverse pregnancy outcomes. While oxygen-sensitive molecular tracers exist for experimental comparison of tissue oxygen saturation *in vivo* (46, 80), these tools have not been applied to models of diet-induced excess adiposity and dysglycemia during pregnancy. Thus, we set out to examine the impacts of maternal excess adiposity and dysglycemia on placental oxygenation and the consequences for the vascular interface adaptations in late gestation. We hypothesized that a state of excess adiposity and dysglycemia arising from chronic preconception high-fat, high-sucrose feeding would lead to placental hypoxia within the labyrinth zone, initiating angiogenic expansion and reduced placental vascular maturity and an altered fetal circulating growth factor milieu in late pregnancy.

## METHODS

### Animal model and sample collection

All animal procedures for this study were approved by the McMaster University Animal Research Ethics Board (Animal Utilization Protocol 20-07-27) in accordance with the guidelines of the Canadian Council on Animal Care. Eight-week-old C57BL/6J female mice (Jackson Laboratories strain 000664) were fed either control rodent chow (CON; 17% fat, 54% carbohydrate, 29% protein (kCal/g), Teklad 22/5 Rodent Diet, Envigo cat. 8640) or high-fat, high-sucrose (HFHS; 45% fat, 17% sucrose, 18% carbohydrate, 20% protein (kCal/g), Research Diets D12451) diet *ad libitum* for a minimum of 12 weeks to model conditions of preconception overweight and obesity, as previously described (19, 37, 81). A comparison of diet compositions is outlined in <u>Supplementary Table 1</u>. At baseline and following 10-weeks following diet onset, female mice were fasted for six hours and subject to oral glucose tolerance tests to assess glycemic control (details below). Following 12- to 15-weeks of dietary intervention, females were time-mated with CON-fed C57BL/6J males to generate pregnancies. Once pregnant, dams were singly housed and allowed to carry pregnancies to embryonic day (E) 17.5. On E17.5, dams were fasted overnight for five hours (03:00 – 08:00 hrs) and administered Pimonidazole HCl (15 mg/ml in isotonic saline; Hypoxyprobe^TM^) intravenously by tail vein injection (60mg/kg of body weight). Successful injection was determined by a lack of resistance and bubbling at the site of injection and appearance of blood following needle withdrawal. Only dams determined to have a successful injection (CON n = 5, HFHS n = 10) were subsequently used for measures of placental hypoxia (details below). Following one hour of pimonidazole administration, blood was sampled for blood glucose measurement and serum collection (see below), and dams were immediately euthanized by cervical dislocation for collection of maternal and fetoplacental tissues.

Uteroplacental tissues were immediately isolated following laparotomy, and the number and position of fetuses and resorptions were recorded. Each fetus was weighed and decapitated, and trunk blood was collected for glucose measures and serum collection. Fetal membranes were collected and snap-frozen for determination of fetal sex by genotyping for the sex-linked *Sry* locus (details below). Placental tissues were collected in an alternating fashion, beginning from the ovarian end of the right uterine horn, being immediately isolated and fixed by immersion in 4% paraformaldehyde in phosphate-buffered saline pH 7.4 for 24 hours at 4°C and processed for paraffin embedding and histology, as previously described (82). Intervening placentas were microdissected on ice to separate labyrinth and junctional zone tissues, using established protocols (83, 84), snap frozen in liquid nitrogen, and stored at -80°C until further processing.

### Measures of glycemic control

Following a six-hour fast (03:00 – 09:00 hrs), blood was sampled from the tail vein for measurement of fasting glucose using a handheld glucometer (Accu-Check Aviva, Roche Diagnostics). 150 μl of blood was collected in heparinized micro-hematocrit tubes (Fisher Scientific) to isolate serum for the measurement of fasting insulin levels before (10 weeks of diet) and during pregnancy (E17.5). Serum insulin concentrations were determined using a commercial ultra-sensitive ELISA (Toronto Biosciences cat. #32380) run according to the manufacturer’s instructions. Samples with absorbance at the background level were set to half of the lower limit of detection (0.0125 ng/ml). At the 10-week time point, mice received an oral gavage of 30% w/v solution of D-glucose in isotonic saline at a dose of 2g/kg of body weight, 30 minutes following initial blood collection. Blood glucose was recorded at 0-, 15-, 30-, 60-, 90-, and 120-minutes after gavage. Preconception glucose handling was measured by the incremental area under the curve (iAUC), using the fasting blood glucose (time 0) as the baseline from which glucose disposal was measured in each animal (64).

### Genotyping for sex determination

Fetal sex was identified by duplex PCR-genotyping of fetal membranes for the Y-linked *Sry* (male sex) and *Fabp2* (autosomal amplification control) loci using primers designed by the Mouse Genetics Core at Washing University School of Medicine at St. Louis (https://mousegeneticscore.wustl.edu/items/pcr-genotyping-primer-pairs/, <u>Supplementary Table 2</u>). Genomic DNA (gDNA) was extracted from snap-frozen fetal membranes using a DNA FastExtract Kit (Wisent Biosciences) according to the manufacturer’s instructions. Amplification products were separated by electrophoresis and visualized under UV illumination using SYBRsafe (ThermoFisher Scientific).

### Serum cytokine and growth factor profiling

In both maternal and fetal serum at E17.5, 11 cytokines and growth factors with known roles in angiogenesis, inflammation and placental development were measured using the Milliplex Mouse Angiogenesis/Growth Factor Magnetic Bead Panel (MAGPMAG-24K, Millipore Sigma), and included the following analytes: Leptin, EGF, IL-6, Endoglin (ENG), Endothelin-1, HGF, Placental growth factor (PlGF), CXCL1, CCL2, Prolactin, and TNF. Samples were thawed, clarified by centrifugation at 12,000×g for 15 minutes and run according to the manufacturer’s instructions. Data were acquired using a Luminex MAGPIX-xMAP system. Following data acquisition, each sample was inspected for quality control. Any sample with an acquisition of fewer than 30 beads was excluded from downstream analysis.

### Tissue processing

Formalin-fixed, paraffin-embedded placentas bisected along the mid-sagittal plane were subjected to systematic serial sectioning using a rotary microtome at a thickness of 5 µm. Sections were collected at 300 µm intervals for morphological assessment using standard hematoxylin and eosin (H&E) staining. Intervening full-thickness sections between a depth between 300 and 750 µm from the cut face were used for analysis. Where possible, paired male and female samples within each litter were used. For all experiments, three replicate sections per sample, each at 75 µm apart, were stained and analyzed. For each sample, composite images of the full tissue section were acquired by scanning and stitching 20× magnification fields using a Nikon NiE Eclipse microscope and NIS Elements software (v5.20.02).

### Immunostaining

For all immunostaining experiments, tissue sections were deparaffinized by incubating at 60°C for 20 minutes, washing with Histo-Clear (National Diagnostics) and rehydrating through graded ethanol solutions to distilled water. Target-specific antigen retrieval conditions are detailed in <u>Supplementary Table 3</u>. Following antigen retrieval, samples were blocked at room temperature with 10% normal goat serum in PBS with 0.1% Tween20 (PBS-T) for one hour. Primary antibodies or lectins were diluted in blocking buffer and applied to tissue sections overnight at 4°C in a humidified chamber. The following day, samples were washed and incubated with fluorophore- or biotin-conjugated secondary detection reagents in PBS-T for 1 hour at room temperature. Antigens requiring tertiary signal amplification (F4/80), sections were washed and incubated for one hour with fluorophore-conjugated Streptavidin diluted in PBS-T. Samples were counterstained with DAPI (Molecular Probes) and mounted using Prolong Gold anti-fade mounting medium (Invitrogen). Positive staining was identified based on comparisons to sections stained with an isotype-matched non-specific primary antibody or no lectin control.

### Quantification of placental hypoxia

To compare oxygen saturation between placentas from CON and HFHS pregnancies, we performed immunofluorescence staining of placental tissue sections using an anti-pimonidazole adduct antibody (PAb27; Hypoxyprobe^TM^). Positive staining was detected using binary thresholding within manually identified regions of interest (ROIs) restricted to each placental zone, using co-immunostaining for SLC16A1 (MCT-1), an established marker for syncytiotrophoblast layer 1 in the murine placenta (85), to define layer boundaries. Thresholds for positive staining were identified based on signal intensity using placental sections stained using the same fluorescent secondary antibody with PAb27 replaced by a titre-matched isotype control. Mean pixel intensity within the binary threshold was measured and recorded using NIS Elements Analysis software and used as a measure of placental oxygen saturation (i.e., higher staining intensity = less oxygen).

### Placental morphometry

Absolute cross-sectional areas of the total placenta and its constituent zones were measured using hematoxylin and eosin (H&E)-stained mid-sagittal sections from each placental sample (CON n = 5 – 6/sex and HFHS n = 8/sex). The perimeter of each layer was traced using NIS Elements image analysis software (Nikon v.5.20.02) based on established morphological criteria (83, 86), and the areas recorded.

### Assessment of placental vascular area and integrity

The proportional area of the placental labyrinth occupied by fetal vasculature was measured using an automated image analysis pipeline constructed using Nikon NIS Elements Analysis software. A visual schematic of pipeline organization and its parameters are outlined in <u>Supplementary Figure 1a</u>. Placentas were fluorescently co-stained for labyrinth pericytes using a monoclonal anti-alpha smooth muscle actin antibody (α-SMA; clone 1A4, Abcam), and the vascular basement membrane using Isolectin β4 from *Griffonia Simplicifolia* (GS-Iβ4, Vector Laboratories) to identify fetal vascular structures within the placenta (15, 86). The region containing GS-Iβ4 staining was used to define the Labyrinth-Junctional Zone boarder which was manually traced to positively identify an ROI for analysis. Images were pre-processed using the *Local Contrast* tool to correct illumination differences across the stitched imaging fields. Positive staining for each marker was separately detected using the *Homogenous Area* function. The resulting binary threshold layers were smoothed at their edges and filtered based on size to remove small clusters of positive pixels (<1µm) not considered to be relevant signals. From this, a singular binary layer (‘Total Vascular Area’; TVA) was generated by combining the two binary layers (<u>Supplementary Figure 1b</u>). This layer was used to refine the original α-SMA^+^ binary to remove pixels detected that were not associated with vascular structures (<u>Supplementary Figure 1c</u>). The area fraction of the TVA binary layer was calculated relative to the manually defined ROI in each image as a surrogate measure of placental vascular abundance. The proportional area of the TVA occupied by α-SMA staining was measured and used as an indicator of microvascular stability, as previously described (87).

### Measures of placental calcification and tissue pathology

Calcification of the placental labyrinth in CON and HFHS placental tissues was assessed by staining placental sections with Alizarin Red S using a modification of previously published protocols (64). Following deparaffinization, tissue sections were incubated in a 1% w/v aqueous solution of Alizarin Red S (Millipore Sigma A5533) with adjusted pH of 6.4 using 10% v/v ammonium hydroxide for 4 minutes. Slides were successively washed in Acetone and 1:1 Acetone/Xylenes (20 dips each), dehydrated by three washes in Xylenes, mounted and coverslipped using Permount^TM^ mounting media (Fisher Scientific). The number of samples with the presence (at least one replicate with positive staining) or absence of staining was recorded. Using NIS Elements Analysis Software (Nikon v5.20.02), the perimeter of the placental labyrinth and chorionic plate was traced to define a region of interest (ROI) for staining quantification. Positive staining was identified using the binary thresholding tool and the total positive area within the ROI was divided by the total ROI area to measure the percent positive area.

In paired serial sections, we also assessed regions coincident with calcium deposition for the presence of placental endothelial cells using fluorescent co-staining with endothelial marker CD31 in the same sections used for quantification of placental vascular areas and integrity (detailed above). In a separate serial section, we also performed staining for SLC16A1 and F4/80 to examine the morphology of the SynT1 layer and placental macrophage infiltration in these regions, respectively. Regions in paired immunostained slides were qualitatively evaluated for the presence or absence of endothelial and trophoblast tissue and assessment of placental macrophage density relative to surrounding, uncalcified regions. Observations were made in duplicate sections for each sample.

### Protein Analyses

#### Western blotting for HIF-1α

Total placental protein was extracted from layer-enriched labyrinth tissue described above. Halves of each labyrinth sample were homogenized in 500 µl of RIPA lysis buffer (50 mM Tris-base, 150 mM NaCl pH 8.0, 0.5% w/v Sodium Deoxycholate, 0.1% w/v Sodium Dodecyl Sulfate, 1% v/v Triton X-100) supplemented with EDTA-free Pierce Protease and Phosphatase Inhibitor Mini Tables (ThermoFisher Scientific) using a FastPrep-24 bead mill (MP Biomedicals) with ceramic beads (4.5 m/s for 30 seconds). Crude lysates were incubated for 30 minutes at 4°C with constant agitation and clarified by centrifugation at 12,000×g for 15 minutes. Protein concentrations were determined using the Pierce^TM^ BCA Protein Assay Kit (ThermoFisher Scientific) and normalized to ∼1mg/ml of protein per sample. 15 µg of total labyrinth protein was separated using 10% separating gel with SDS-PAGE and transferred to PVDF membranes using a TransBlot Turbo Transfer System (1.0 A 25V for 28 min; 1704150, BioRad). Membranes were blocked for one hour at room temperature in Tris-buffered saline supplemented with 1% Triton X-100 and 5% skim milk powder, cut based on target molecular size, and incubated overnight with either rabbit anti-HIF-1α (NB100-479, Novus Biologicals; 1:1000) or HRP-conjugated rabbit-anti-β-actin (Cell Signalling Technology 5125S; 1:15,000). Blots for HIF-1α were further incubated with an anti-rabbit IgG secondary antibody (ab6721, Abcam; 1:100,000). Proteins of interest were detected using Clarity Max Western ECL Blotting Substrate (1705062, BioRad) and images were developed using a ChemiDoc MP Imaging System (1708280, BioRad). Densitometric quantification of images was performed using ImageLab software (BioRad). HIF-1α signal intensity was normalized to that of β-actin to determine relative expression.

#### Protein array

A Mouse XL Cytokine Proteome Profiler Array (ARY028, R&D Systems) was used to profile the expression of 111 proteins in placental labyrinth zone lysates. Labyrinth tissues were prepared as above, with the exception that tissues were extracted in a non-denaturing lysate extraction buffer (50 mM Tris-base, 150 mM NaCl, 20 mM NaF) supplemented with Pierce Protease Inhibitor Cocktail (ThermoFisher Scientific) containing Aprotinin, Bestatin, E-64, Leupeptin, AEBSF, Pepstatin A, and EDTA. Following homogenization, an extraction buffer containing 10% Triton X-100 was added to a final concentration of 1%. Lysates were incubated at 4°C for 30 minutes and clarified by centrifugation. Lysates were pooled according to maternal diet and placental sex (n=8 – 11/group) to a total of 200 µg of protein with equal contribution from each sample and applied to array membranes, according to the manufacturer’s instructions. Signal densitometry was measured using the ‘Volume’ tool with global background subtraction in ImageLab software (Bio-Rad, https://www.bio-rad.com/en-ca/product/image-lab-software) Semi-quantitative differences in protein expression were measured by comparing the fold difference of HFHS samples to CON samples within each sex. A complete list of analytes, membrane coordinates, and their signal intensities for each membrane are available in <u>Supplementary Table 4</u>.

### Statistical Analysis

For all analyses in this study, one litter or dam was considered a biological replicate, with one male and female fetus or placenta from each litter used where data are stratified by sex. For all pairwise comparisons between CON and HFHS dams, data were analyzed using Welch’s t-test or Mann-Whitney U-test where the data were discrete values (e.g., litter size) or non-normally distributed, as determined by D’Agostino-Pearson K^2^ test. For time course experiments, data were analyzed using repeated two-way ANOVA based on general linear modelling with Geisser-Greenhouse correction, with maternal diet and time as independent variables. Where a significant main effect or interaction was found, pairwise post-hoc comparisons were made with correction for multiple comparisons using Bonferroni’s method. Data were analyzed in Prism 9 (v.9.5.1, GraphPad Software, La Jolla, California, USA, www.graphpad.com). For all quantitative fetal and placental analyses comparing CON and HFHS groups with replicate measures, data were analyzed using mixed effects linear models using the lmerTest package (88) (version 3.1-3, SCR_015656) in R (v.4.2.3) with maternal diet and fetoplacental sex as fixed effects, and sample as a random effect to nest repeated measurements. Main effects of diet and sex were measured in these models using two-way ANOVA with Type 3 sum of squares and Satterthwaite’s method for calculating denominator degrees of freedom. Odds ratios of categorical fetal weight outcomes (small and large for gestational age), were calculated using nested binary logistic regressions based on generalized linear mixed effects modelling (glmer) using the lme4 package (version 1.1-33, RRID: SCR_015654) (89), with diet and sex as fixed effects and dam as a random effect to nest repeated measures within a litter. Where significant main effects of interactions were present in linear models, pairwise post-hoc comparisons were made between the estimated marginal means using the emmeans package (90) (v.1.8.5, RRID SCR_018734). Model summary tables (<u>Supplementary Tables 5</u> <u>and 6</u>) were generated using the sjPlot package (91) (v.2.8.14). Correlations between datasets were performed using Pearson correlation in R For all analyses, p<0.05 was used as a nominal threshold for determining statistical significance.

## RESULTS

### High-fat, high-sucrose diet feeding induces preconception excess adiposity and dysglycemia

Female mice were fed a control rodent chow (CON), or an obesogenic diet high in saturated fat and sucrose (HFHS) for a minimum of 12 weeks before mating to model a state of maternal preconception excess adiposity, as previously described (19, 37, 39, 81, 92). Female mice fed an HFHS diet were significantly heavier than CON-fed females from six weeks of diet onset (<u>Figure 1b</u>), and exhibited moderately elevated fasting blood glucose, hyperinsulinemia and insulin resistance (HOMA-IR) by 10 weeks of feeding (<u>Figure 1c–e</u>). Despite these glycemic impairments, HFHS females were not glucose intolerant on an oral glucose tolerance test (p = 0.302), although they did show an altered pattern of glucose clearance (time × diet interaction, p = 0.026, <u>Figure 1f</u>) and remained hyperglycemic throughout the test compared to CON (main effect of diet p = 0.013; <u>Supplementary Figure 2a</u>). Glucose tolerance was not correlated with the degree of weight gain in either group (p ≥ 0.90; <u>Supplementary Figure 2b</u>).

**Figure 1.**
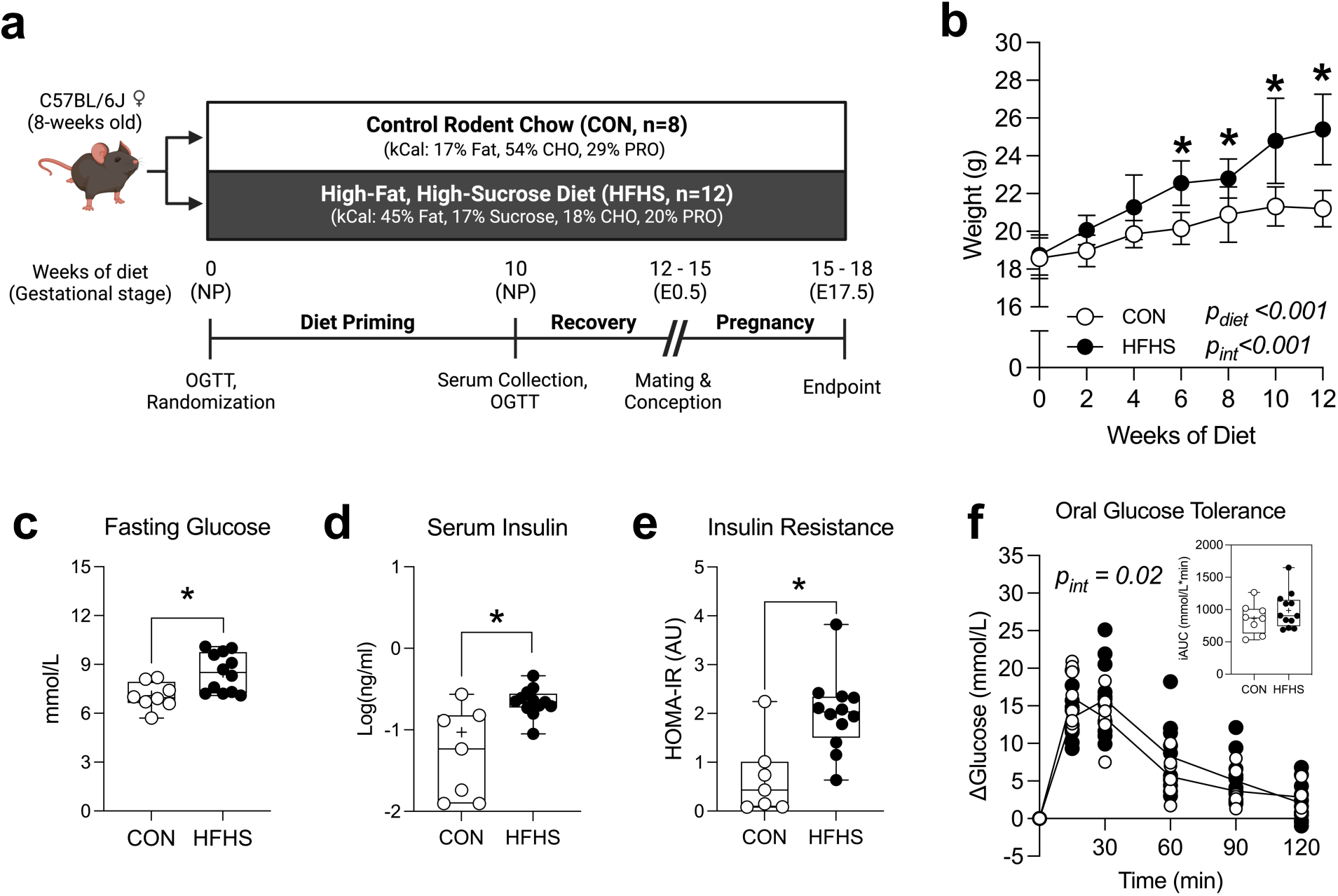
High-fat, high-sucrose diet induces excess weight gain, hyperglycemia, and insulin resistance prior to conception. **(a)** Experimental timeline; pregnant dams were euthanized at embryonic day 17.5 (E17.5) following 12-15 weeks of preconception dietary intervention. **(b)** Pre-pregnancy female body weights during 12 weeks of dietary intervention. **(c)** Six-hour fasted blood glucose and **(d)** serum insulin levels after ten weeks of diet intervention. **(e)** Homeostatic model assessment of insulin resistance (HOMA-IR). **(f)** Changes to blood glucose levels following oral glucose challenge (2g/kg body weight) after 10-weeks of diet intervention; inset graph – area under the glucose curve. Data for CON (n= 7 – 8) and HFHS (n=12) are presented as mean ± standard deviation (SD) for (b). Boxplots in (c – f) display min and max values (whiskers), interquartile range (IQR, box) and median (line), with ‘+’ denoting the group mean. Individual data points from each repeated measurement over time are displayed in (f). Data were analyzed by repeated two-way ANOVA with post-hoc Bonferroni-corrected pairwise comparisons (b, f), or by pairwise comparison with Welch’s t-test or Mann-Whitney U-test (c – e). P-values from two-way ANOVA are indicated where main effects were statistically significant. *Indicates p<0.05 for pairwise comparisons. CON = control diet (open circles), HFHS = high-fat, high sucrose diet (closed circles). kCal = proportion of kilocalories from each macronutrient source; CHO = carbohydrates; PRO = protein; NP = non-pregnant; E0.5 = embryonic day 0.5; OGTT = oral glucose tolerance test; HOMA-IR = homeostatic model assessment of insulin resistance.

### HFHS diet feeding modifies metabolic adaptations to pregnancy

CON and HFHS females were time-mated and conceived between 12 – 15 weeks of diet onset. HFHS females exhibited similar mating efficiency, fertility, and rates of fetal loss compared to controls (<u>Table 1</u>). On average, HFHS females gained more than double the weight of CON females relative to their baseline body weight and weighed significantly more at the time of conception (E0.5; <u>Figure 2a and b</u>). While HFHS dams remained heavier than CON dams during pregnancy, they exhibited an altered pattern of weight gain (time × diet interaction, p < 0.001; <u>Figure 2b</u>) and gained less overall weight throughout gestation (<u>Figure 2c</u>). This was attributable to changes in maternal rather than fetal growth as total litter weights were similar between diet groups (<u>Table 1</u>). In keeping with altered patterns of maternal weight gain, HFHS dams differed in average daily caloric intake in different periods of pregnancy (time × diet interaction, p < 0.001), with increased daily caloric intake in early gestation (E6.5 – 8.5) but decreased daily intake in late gestation (E14.5 – 17.5, <u>Figure 2d</u>), consistent with previous reports (81, 93, 94). Maternal adiposity was elevated in gonadal but not mesenteric depots (<u>Figure 2e</u>) and was associated with an overall elevation in circulating leptin levels in HFHS dams at E17.5 (<u>Figure 2f</u>).

**Figure 2.**
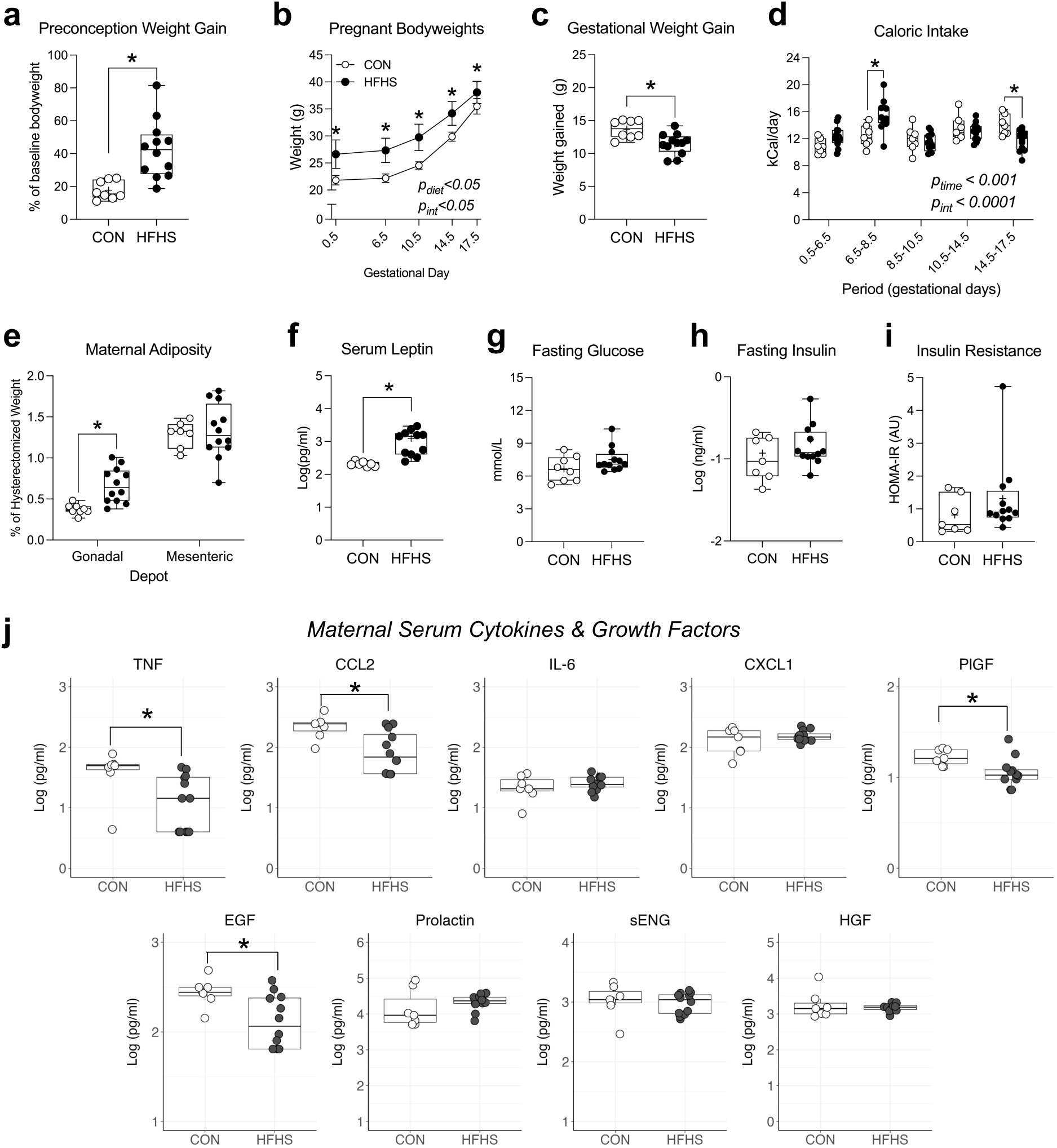
High-fat, high-sucrose feeding modifies maternal adaptations to pregnancy. **(a)** Weight gain relative to baseline body weight at the time of conception. **(b)** Maternal body weight from gestational day 0.5 to 17.5. **(c)** Total weight gained from gestational day 0.5 to 17.5. **(d)** Average calorie intake per day based on total food intake during indicated gestational periods. **(e)** Weights of gonadal and mesenteric adipose depots relative to hysterectomized bodyweight at E17.5. **(f)** Six-hour fasting serum leptin, **(g)** blood glucose, and **(h)** serum insulin levels at E17.5. **(i)** Homeostatic model assessment of insulin resistance (HOMA-IR) at E17.5. **(j)** Multiplex profiling of inflammatory, endocrine, and angiogenic mediators in maternal serum at E17.5 (from left to right, top: Tumor necrosis factor (TNF), chemokine C-C motif ligand 2 (CCL2), interleukin 6 (IL-6), chemokine C-X-C motif ligand 1 (CXCL1), Placental growth factor (PlGF); bottom: Epidermal growth factor (EGF), Prolactin, soluble Endoglin (sENG), Hepatocyte Growth Factor (HGF)). Data for CON (n = 7 – 8) and HFHS (n = 8 – 12) are presented as mean ± SD for (b). Boxplots in (a, c – i) display min and max values (whiskers), IQR (box), and median (line), with ‘+’ denoting the mean value for each group. Boxplots in (j) display min and max values within 1.5 IQR (whiskers), with IQR (box) and median (line). Data in (b, d) were analyzed using repeated measured two-way ANOVA with post-hoc Bonferroni pairwise comparisons; all other data were analyzed by pairwise comparison with Welch’s t-test or Mann-Whitney U-test (a, c, e – j). P-values from two-way ANOVA are indicated where the main effects were statistically significant. *Indicates p<0.05 for pairwise comparisons.

**Table 1.** Measures of mating efficacy and reproductive outcomes at E17.5.

| Measure | CON (n=7 – 8) | HFHS (n=11 – 12) | *p* |
| --- | --- | --- | --- |
| Maternal age (weeks) | 21.4<br>(20.3 – 22.3) | 21.5<br>(20.3 – 22.9) | 0.73<sup>b</sup> |
| Pairings to 1<sup>st</sup> plug | 1.50<br>(1 – 3) | 1.42<br>(1 – 3) | 0.90<sup>a</sup> |
| Plugs to pregnancy | 1.00<br>(1) | 1.08<br>(1 – 2) | 0.99<sup>a</sup> |
| Implantation sites | 8.63<br>(8 – 9) | 8.83<br>(7 – 11) | 0.99<sup>a</sup> |
| Resorption rate (%) | 8.85<br>(0 – 25) | 10.58<br>(0 – 45) | 0.95<sup>a</sup> |
| Viable fetuses | 7.88<br>(6 – 9) | 7.75<br>(6 – 9) | 0.79<sup>a</sup> |
| % Female | 47.84<br>(33 – 85) | 57.67<br>(29 – 83) | 0.28<sup>a</sup> |
| Total Litter Weight | 5.91<br>(5.36 – 6.76) | 5.77<br>(4.83 – 6.51) | 0.67<sup>b</sup> |

Data are presented as mean with range (min – max). Data were analyzed using <sup>a</sup>Mann-Whitney U-test or <sup>b</sup>Welch's t-test.

Preconception metabolic impairments were diminished by E17.5 with similar levels in maternal fasting blood glucose (<u>Figure 2g</u>), fasting serum insulin levels (<u>Figure 2h</u>), and HOMA-IR (<u>Figure 2i</u>) between CON and HFHS dams, consistent with previous reports in similar dietary models and obese humans (15, 30, 94–99). These data demonstrate that late gestational adaptations to lipid and glucose handling differ between CON and HFHS pregnancies. Normalization of some maternal metabolic indices could indicate changes to macronutrient mobilization to the fetoplacental unit in late pregnancy.

Excess adiposity and metabolic impairments during pregnancy have previously been associated with increased circulating inflammatory cytokines and growth factors which may act on placental and fetal tissues (15, 100, 101). Therefore, we profiled levels of maternal circulating cytokines, chemokines, and growth factors previously identified to be altered in pregnancies complicated by maternal obesity and with known roles in vascular and placental function (100, 102). Unexpectedly, but in keeping with our observations of a relatively ‘improved’ metabolic phenotype in late gestation, we observed *reduced* levels of circulating pro-inflammatory mediators TNF and CCL2 in maternal serum at E17.5. In addition, we also observed decreased circulating levels of cytokines including EGF and PlGF. Levels of IL-6, CXCL1, prolactin, sENG, and HGF were not different between groups (<u>Figure 2j</u>). These data suggest a state of diminished endocrine and inflammatory adaptations in HFHS dams that are normally observed with the progression of pregnancy, similar to some recent reports in human cohorts (30, 99, 100, 103–105).

### HFHS diet-induced excess adiposity is associated with placental hypoxia

To determine whether HFHS diet-induced excess adiposity impaired placental oxygenation in late gestation (E17.5), we compared global pimonidazole adduct formation as a readout of tissue hypoxia in each placental layer. We found that both the junctional and labyrinth zones of HFHS placentas were hypoxic compared to CON (<u>Figure 3a – b</u>). In all samples, the junctional zone showed a greater signal intensity for hypoxic adduct formation, consistent with the known perfusion pattern of the murine placenta (72). The reduction of oxygen availability in both tissue layers (as opposed to solely the junctional zone), suggests that hypoxia arises, in part, from insufficient oxygen supply, rather than from increased fetoplacental oxygen extraction and thus reduced oxygen supply to junctional zone tissue. To determine whether reductions in tissue oxygenation were sensed as acute hypoxic stress within the labyrinth, we measured HIF-1α protein levels in layer-enriched labyrinth tissue samples. Relative levels of intact (120 kDa) HIF-1α were similar between CON and HFHS placentas (p = 0.9779, <u>Figure 3c</u>), suggesting that while placental oxygen levels are depleted in HFHS pregnancies, this did not activate a global acute hypoxic stress response.

**Figure 3.**
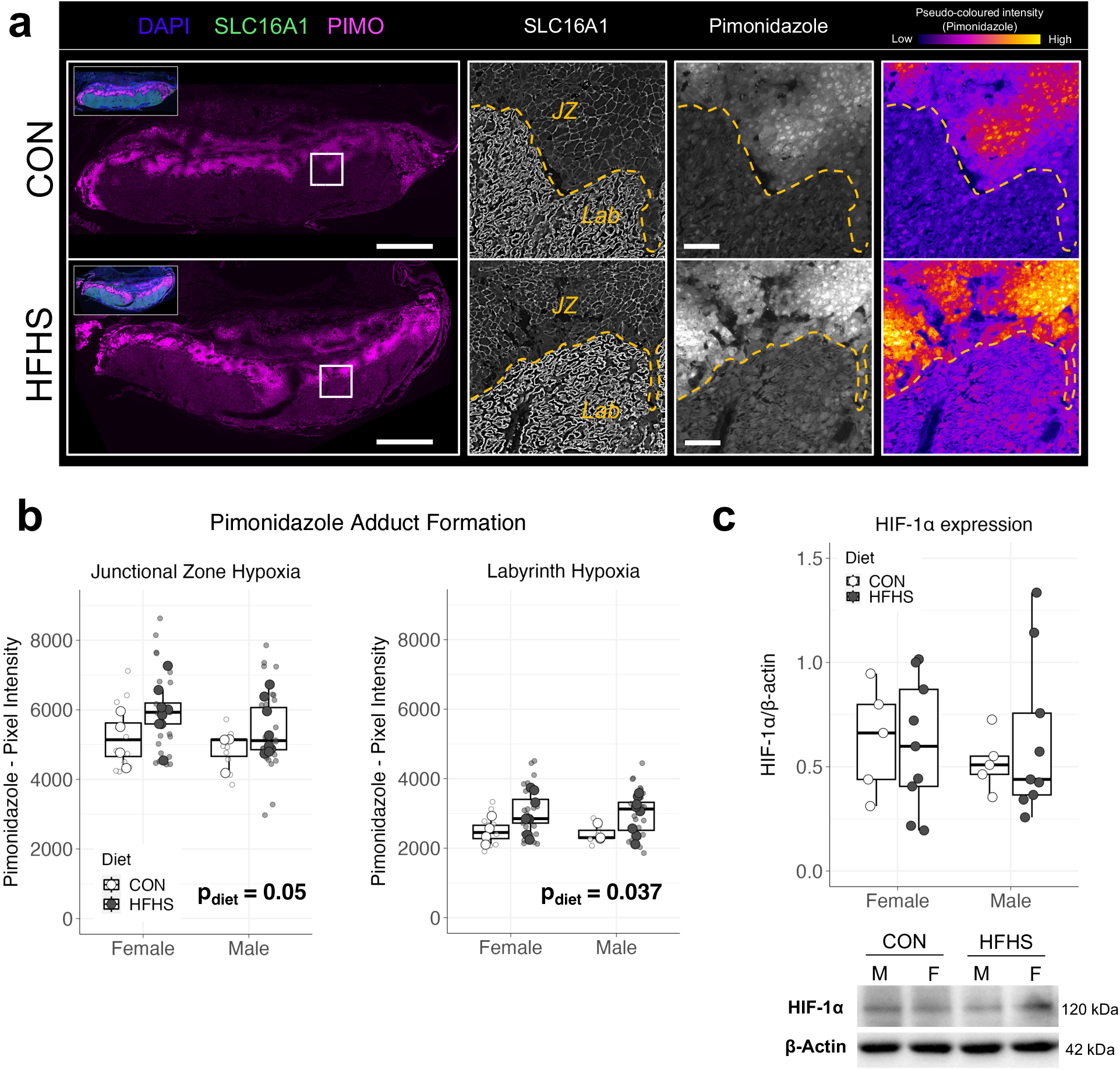
HFHS diet-induced excess adiposity is linked to placental hypoxia. **(a)** Representative staining of CON and HFHS placentas at E17.5 for pimonidazole adduct formation; left –widefield composite images of anti-pimonidazole (PIMO, magenta) immunostaining, right inset shows multichannel overlay of nuclear counterstaining (DAPI, blue), SLC16A1 (monocarboxylate transporter 1 (MCT-1), green) and Pimonidazole protein adducts (PIMO). Scale bar = 1000 µm. White box inset defines location of fields in panels to the right. Middle-left panel – SLC16A1 staining used to define boundaries (hashed line) of Labyrinth (Lab) and Junctional Zones (JZ) defined for image analysis. Middle-right panel – staining of pimonidazole adducts in the JZ and labyrinth used for measurements of hypoxia based on staining intensity of pimonidazole adducts. Right panel – pimonidazole immunostaining pseudo coloured according to pixel intensity. **(b)** Quantification of pimonidazole immunostaining fluorescence signal according to the mean pixel intensity within the Junctional Zone and Labyrinth compartments stratified by fetoplacental sex in CON (n = 3 – 4/sex) and HFHS (n = 8/sex) placentas. **(c)** HIF-1α protein expression measured using Western blot in layer-enriched labyrinth tissue from CON (n = 5/sex) and HFHS (n = 8 – 9/sex) placentas. Protein expression was quantified using densitometry and normalized to the expression of β-Actin (loading control) derived from the same membrane. Boxplots in (b) depict the min and max values within 1.5 IQR for each group (whiskers), IQR (box), and median (line). Large data points depict the mean value for each sample analysed in triplicate, small data points depict individual mean intensity of each sample used for analysis. Boxplots in (c) depict min and max values (whiskers) with IQR (boxes) and median (line). Data were analysed using two-way ANOVA on mixed effects linear models with diet and sex as fixed effects and sample (placenta) as a random effect in (b) or regular two-way ANOVA in (c). P-values from two-way ANOVA are indicated where main effects were statistically significant. kDa = kilodalton.

### Gross placental morphology is not markedly altered in HFHS-fed pregnancies

Placental hypoxia has been associated with diet-induced excess adiposity and metabolic impairments and with changes to placental microvascular abundance and stability. However, the reported direction of these changes varies between models (15, 19, 63, 65). We and others have shown that preconception diet-induced obesity increased microvascular area within the placenta with concomitant reductions in the stability of these vessels, as measured by a reduction of perivascular cells (α-SMA^+^ pericytes) relative to the increased vascular endothelial cells, and that this may be driven by placental adaptations to chronic hypoxia (15–17, 19). To further qualify the degree of placental labyrinth zone hypoxia, we performed gross morphometric analyses for placental vascular structures. Mid-sagittal sections of HFHS and CON placentas had similar cross-sectional total placental areas (<u>Figure 4a</u>) and no group effect on the proportional areas of either the labyrinth or junctional zones (<u>Figure 4b and c</u>). There was a tendency for males to have larger placentas (main effect of sex, p = 0.078), driven by a significant increase in the cross-sectional area of the junctional zone (<u>Figure 4b</u>). We also observed a specific reduction of junctional zone area in HFHS female fetuses compared to HFHS males (p = 0.025). We next examined whether HFHS induced specific changes to placental microvascular abundance or stability within the labyrinth via co-staining of placental sections with GS-Iβ4 lectin and immunostaining for α-SMA to mark vascular membranes and perivascular cells, respectively (<u>Figure 4d</u>). No differences in total placental vasculature were noted between the groups (<u>Figure 4e</u>). Similar ratios between total vessel area and α-SMA^+^ pericytes were observed between CON and HFHS placentas, suggesting placental vascular network stability and a lack of neoangiogenic response to hypoxia. These data suggest that the reduced oxygenation of the placental labyrinth in HFHS pregnancies is not representative of acute hypoxic stress, which is consistent with similar HIF-1α expression observed between groups.

**Figure 4.**
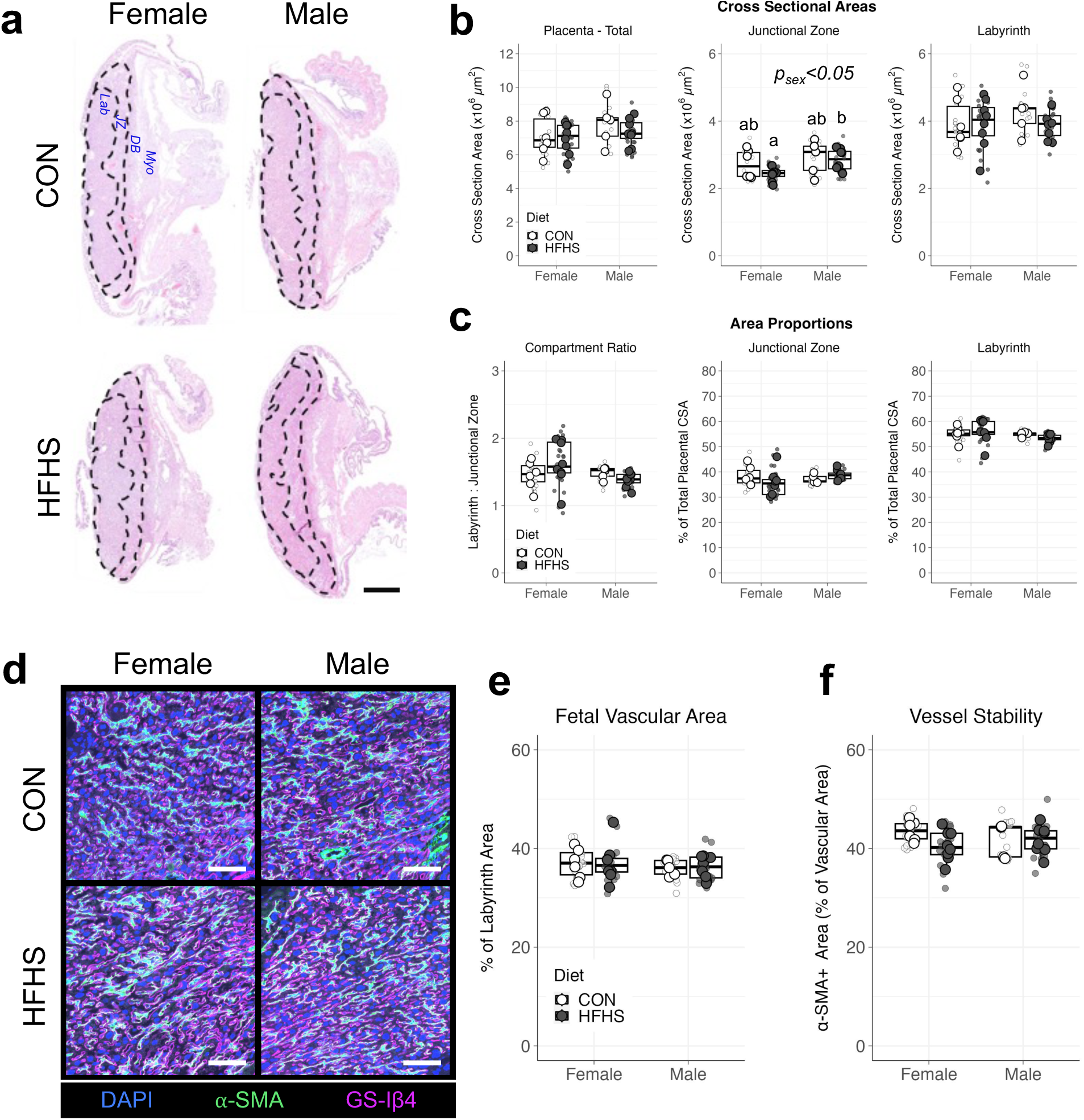
Gross placental morphology and microvascular stability are similar between CON and HFHS pregnancies. **(a)** Representative composite images of H&E-stained placentas. Boundaries between Labyrinth (Lab), Junctional Zone (JZ), and maternal tissue (decidual basalis (DB) and myometrium (Myo)) were defined based on tissue architecture (hashed lines); scale bar = 1000 µm. **(b)** Cross sectional areas of the total placenta (combined JZ and Lab), individual compartments, **(c)** area ratios, and proportional areas were measured for placentas from CON and HFHS pregnancies. **(d)** Representative multi-channel overlay of fields from the Labyrinth of CON and HFHS placentas co-stained for the vascular basement membrane and perivascular cells (pericytes) using isolectin beta 4 from *Griffonia Simplicifolia* (GS-Iβ4, magenta), anti-alpha smooth muscle actin monoclonal antibody (α-SMA, green), respectively; scale bar = 100 µm. **(e)** Proportional area within the Labyrinth occupied by fetoplacental vasculature, defined using combined staining of GS-Iβ4 and α-SMA (Fetal Vascular Area). **(f)** Stability of fetoplacental vasculature measured according to the percent of the Fetal Vascular Area occupied by α-SMA positive staining Data are presented with boxplots depicting the min and max values within 1.5 IQR (whiskers), IQR (box), and median (center line). Large data points represent the mean of replicate measures from sample replicates, small data points depict individual values of each sample replicate. Data were analysed using two-way ANOVA on mixed effects linear models with diet and sex as fixed effects and sample (placenta) as a random effect. P-values are indicated where main effects were statistically significant. Boxplots without shared letters are significantly different from one another according to post-hoc comparison of estimated marginal means. All samples were analysed in triplicate for CON (n = 5 – 6/sex) and HFHS (n = 8/sex) placentas. All samples are taken from composite images imaged at 20× magnification.

### The labyrinth proteome in HFHS placentas is altered in a sex-dependent manner

In addition to hypoxia, placental inflammation and oxidative stress have also been shown to perturb fetoplacental vascular homeostasis in pregnancies complicated by excess adiposity (27, 106). Therefore, we next investigated these and related signalling pathways in the placenta and hypothesized that these pathways may underpin the relative placental hypoxia in HFHS pregnancies. Tissue lysates were generated from layer-enriched placental labyrinth tissues and pooled by diet group and placental sex (CON n = 7 – 8/sex and HFHS n = 10 – 11/sex). Semi-quantitative grouped expression of 111 protein analytes involved metabolism, vascular biology, and inflammatory regulation were probed using spotted membrane protein arrays (Supplementary Table 4). Sixty-nine target proteins were detectable at a level that allowed for reliable comparisons of expression between diet groups (<u>Figure 5a</u>). Thirty-three proteins with altered expression in the placenta of either HFHS females, males, or both sexes were identified using within-sex comparisons and an absolute fold-change of ≥1.25 as a threshold for biologically relevant differences (37). Importantly, notable differences in the expression of select analytes between sexes in each group were identified in pooled protein lysates using this cut-off. This included lower expression of Secreted phosphoprotein 1 (SPP-1) and Osteoprotegerin (OPG; <u>Figure 5b</u>) which are suppressed by androgen signalling in other contexts (107, 108) highlighting the ability of this assay to detect biologically relevant differences.

**Figure 5.**
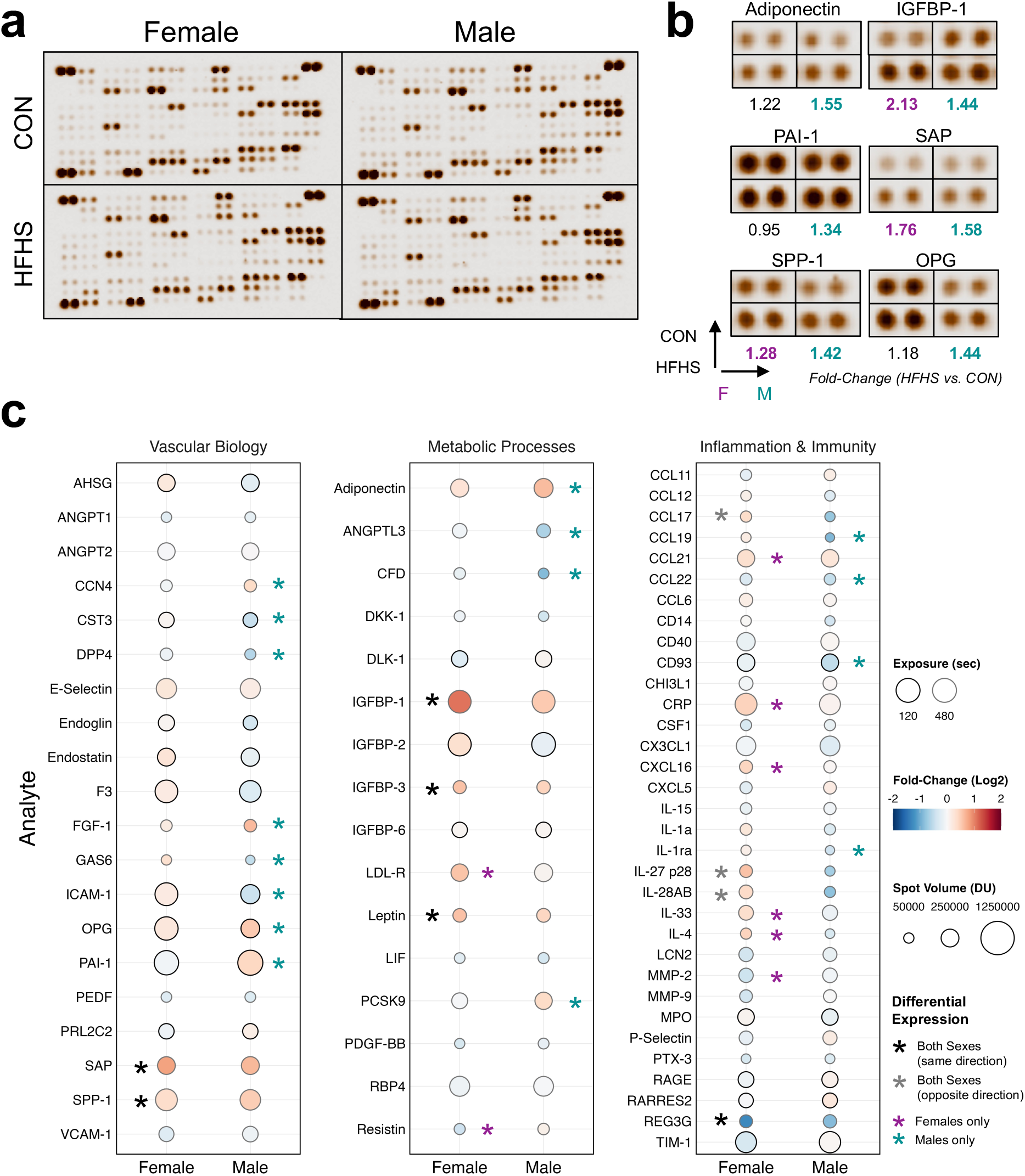
Placental labyrinth expression of mediators related to inflammation and vascular damage are altered in HFHS pregnancies. **(a)** 111 protein analytes were probed in duplicate in pooled labyrinth tissues from CON (n = 8/sex) and HFHS (n = 10 – 11/sex); all analytes and coordinates are listed in Supplementary Table 4. **(b)** Representative duplicate spots of analytes considered to be differentially expressed (≥1.25 absolute fold-change (FC) in spot volume (densitometric units (DU) between HFHS vs. CON) in either sex. Coloured text indicates FC in females (magenta) or males (teal) passing the differential expression cut-off. **(c)** Dot plot of all 69 reliably detected target analytes in the protein array. Target analytes were grouped according to known functions and gene ontology (GO) annotations into three categories – vascular biology, metabolic processes, and inflammation and immunity. Dot outlines indicate whether measures were taken from membranes at lower (120 seconds, black outline) or higher exposure (480 seconds, grey outline). Dot colour corresponds to the Log_2_(Fold-Change) (HFHS versus CON) for comparisons within each sex. Dot size corresponds to the absolute spot volume at a given exposure. Asterisks indicate target analytes considered to be differentially expressed in both sexes in the same (black) or opposite (grey) direction, or exclusively in females (magenta) or males (teal). IGFBP-1 = insulin-like growth factor binding protein 1; PAI-1 = Plasminogen activator inhibitor 1; SAP = Serum amyloid P component; SPP-1 = Secreted phosphoprotein 1; OPG = Osteoprotegerin.

HFHS placentas had altered expression of proteins involved in metabolism and lipid handling such as Angiopoietin-like 3 (ANGPTL3) and Proprotein convertase subtilisin/kexin type 9 (PCSK9) in males, and Low-density lipoprotein receptor (LDL-R) in females, with HFHS showing fold changes of -1.27, 1.31, and 1.48, respectively. Both sexes exhibited increases in leptin (female fold change 1.35 and male fold change 1.51). Surprisingly, adiponectin, whose expression is usually decreased in maternal circulation with obesity (109), exhibited increased relative expression in HFHS labyrinth tissue, although only male placentas met our cut-off (male fold change of 1.55, versus female fold change of 1.22; <u>Figure 5b</u>). Both hypoxia and adiponectin signalling are known to positively regulate factors that reduce growth factor availability, including insulin-like growth factor binding proteins (IGFBPs) (110–112) which limit IGF-1 and IGF-2 bioavailability to limit fetal growth (113). Both IGFBP-1 and IGFBP-3 expression were increased in HFHS labyrinth tissue of both sexes (IGFBP-1: male fold change 1.44, female fold change 2.13; IGFBP-3 male fold change 1.36, female fold change 1.50).

Several proteins involved in responses to tissue injury and inflammation were also altered in HFHS labyrinth tissue compared to controls. This included upregulation of members of the acute phase family of proteins Serum amyloid component P in both sexes (SAP; male fold change 1.76, female fold change 1.58) and C-reactive protein in females (CRP, female fold change 1.37). HFHS females had markedly increased levels of cytokines and chemokines related to type 2 immunity and tissue repair (114, 115), including IL-4 and IL-33 (fold change 1.27 and 1.36, respectively). Similarly, chemokines and cytokines related to T_h_2/T_reg_ cell recruitment and induction of IL-10 (116–118) displayed discordant changes, increasing in females but decreasing in males (CCL17; female fold change 1.29, male fold change -1.69 and IL-27 p28; female fold change 1.49, male fold change -1.61). Male-specific increases were observed in mediators of vascular biology including increased Plasminogen activator inhibitor 1 (PAI-1; fold change 1.34) and Fibroblast growth factor 1 (FGF-1; fold change 1.56). Several factors related to fibrosis and vascular tissue calcification(119–122) including SPP-1 in both sexes (male fold change 1.28, female fold change 1.42), and OPG and WNT1-inducible-signaling pathway protein 1 (CCN4) were also altered in males (fold change 1.44 and 1.31, respectively). These data suggest sex-specific altered macronutrient processing, growth factor regulation, and inflammatory homeostasis within the HFHS placental labyrinth in late gestation.

### HFHS placentas demonstrate signs of inflammatory injury, malperfusion, and calcification

The altered expression of signalling factors related to inflammation, fibrosis, and wound repair we observed in HFHS labyrinth lysates prompted a more specific comparison of histopathology between CON and HFHS placentas. We observed a prominent and varied degree of pathological changes between HFHS and CON placentas of both sexes. Among these changes, the most readily identifiable was the deposition of a fibrinoid-type material, a stereotypical sign of maternal placental malperfusion (35, 123). We also observed acellular deposits resembling tissue calcification (<u>Figure 6a</u>). Recent reports using similar models of maternal high-fat diet intervention have identified increases in placental calcification (39, 64). We performed histochemical staining to identify calcium phosphate deposits within the labyrinth (<u>Figure 6b</u>) and confirmed that HFHS placentas show a markedly increased incidence of placental calcification (CON: 1/11 vs. HFHS: 13/16 placentas, p = 0.003). Overall, HFHS placentas had a significantly increased proportional area within the labyrinth occupied by calcium deposits. The strength of this association was stronger in males compared to females when performing pairwise comparisons (male: 0/5 vs. 7/8 samples, female: 1/6 vs. 6/8 samples, CON vs. HFHS, Bonferroni adjusted p = 0.009 and 0.103, respectively; <u>Figure 6b</u>). Of note, the extent to which the labyrinth tissue was calcified in HFHS placenta varied widely, ranging from complete absence to small, localized foci, to widespread and large depositions (<u>Figure 6c</u>); this variation occurred even within siblings from the same pregnancy.

**Figure 6.**
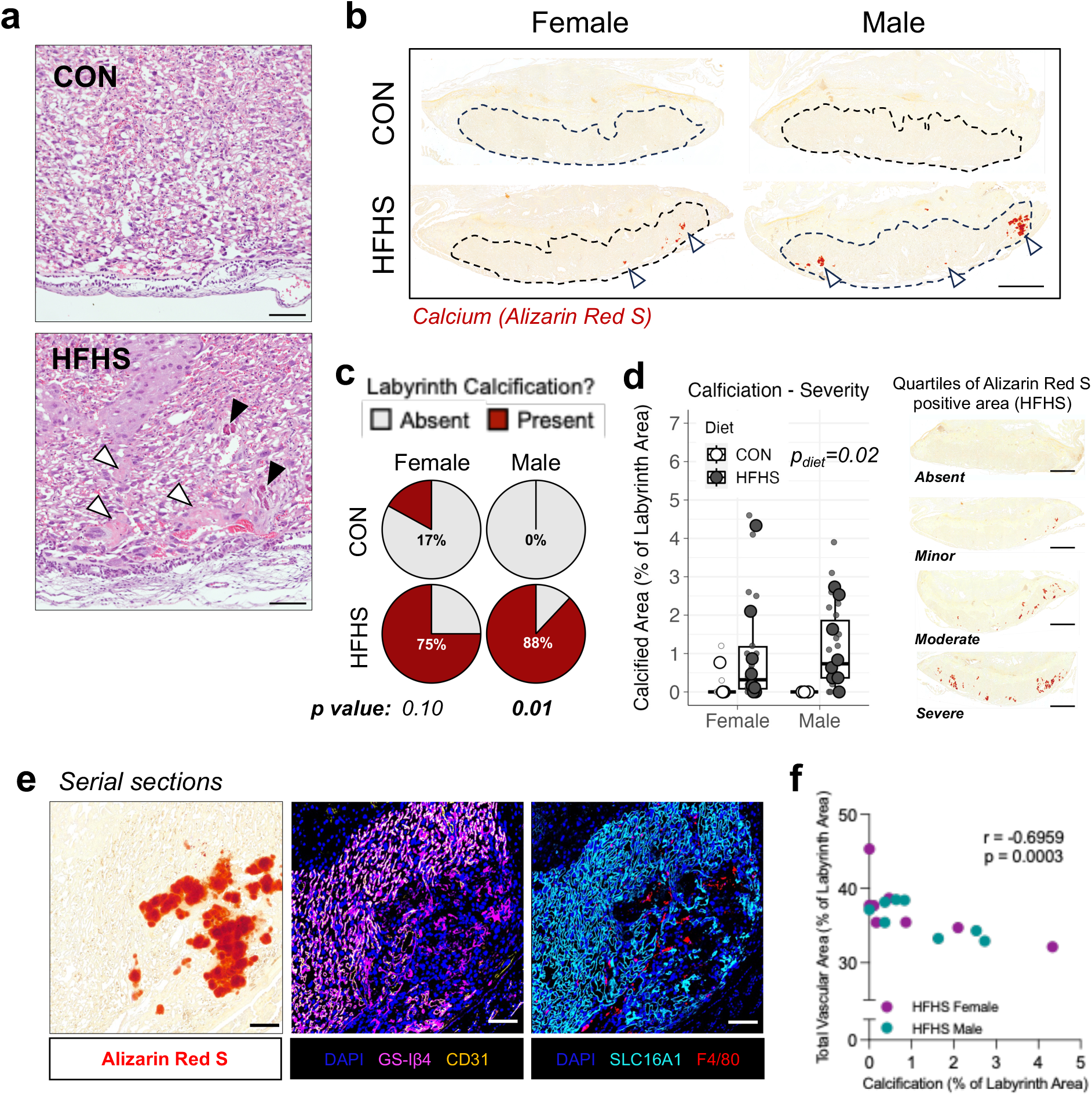
HFHS diet-induced excess adiposity is linked to focal dystrophic calcification and signs of malperfusion damage and inflammation in the placental labyrinth at late gestation. **(a)** H&E staining of CON and HFHS placental tissues depicting representative histopathology of CON and HFHS placentas. The presence of tissue lesions consistent with occlusions/calcification (black arrows) and eosinophilic fibrinoid deposition (white arrows) were more frequently observed in HFHS samples; scale bar = 100 µm. **(b)** Representative composite images of tissue sections from CON (n = 5 – 6/sex) and HFHS (n = 8/sex) stained for calcium deposition using Alizarin Red S. Hashed outlines indicate the labyrinth zone, to which analysis was restricted. White arrows indicate calcified foci; scale bar = 1000 µm. **(c)** Percent of placental samples with the presence of calcium in at least one replicate tissue section (3 replicates/sample). **(d)** Quantification of calcification severity based on the percent Alizarin Red S positive area; images depict representative samples each quartile of Alizarin Red S staining within the HFHS group (both males and females); scale bar = 1000 µm. **(e)** Representative fields of the labyrinth from serial placental sections stained for calcium (left panel), placental vascular endothelium (GS-Iβ4, CD31; middle panel), or syncytiotrophoblast and macrophages (SLC16A1 and F4/80, respectively; right panel); scale bar = 100 µm. **(f)** Correlation of mean placental labyrinth calcification and fetoplacental vascular proportional areas in HFHS female (magenta) and male (teal) samples. Data in (c) are presented with boxplots depicting the IQR (box), median (center line) and min and max values within 1.5 IQR for mean values (whiskers); large data points represent the mean value of triplicate measures per sample, small data points represent individual replicates. Data points in (f) represent the mean value for each sample. All representative image fields were taken from composite stitched images of whole tissue sections imaged at 20× magnification. Data were analysed using Fisher’s exact test with post-hoc Bonferroni correction for multiple testing (b), two-way ANOVA on mixed effects linear models (c) or Spearman correlation (f).

Tissue calcification can arise by several means including active processes of mineralization (metastatic) or tissue injury (dystrophic). The latter includes areas of thrombosis and infarction of both maternal and fetal origin (124–126), as well as inflammatory or infectious stimuli (127). In line with previous reports (64), we observed that areas of labyrinth calcification coincided with an absence of CD31^+^ endothelial cells. In these same areas, residual staining of vascular basement membranes by GS-Iβ4 remained, indicating these were areas once occupied by labyrinth vasculature (<u>Figure 6e</u>). These areas also lacked staining of SLC16A1, indicating an absence of maternally facing syncytiotrophoblast layer 1 (SynT-I) cells. Coincident with this, we also found a focal accumulation of F4/80^+^ cells, indicating macrophage recruitment. In a few rare instances (n=2/16 HFHS samples), we observed calcium deposits in regions outside of the labyrinth zone, affecting the border between the decidual and junctional zone (<u>Supplementary Figure 2</u>) in what appeared to be either maternal arterial or venous channels. While we cannot discern the exact triggers of these changes, our results suggest that these calcified areas are of a dystrophic type, resulting from the destruction of vasculosyncytial membranes. We observed a strong negative correlation between placental vascular area and the proportion of the labyrinth occupied by Alizarin Red S staining (Pearson’s r = -0.696, p = 0.003), indicating that while previous measures of total placental vasculature were not different between CON and HFHS groups, the area available for oxygen and nutrient exchange may be relatively compromised for HFHS fetuses.

### Maternal HFHS diet intake alters circulating fetal inflammatory mediators

Given the evidence of tissue damage affecting fetoplacental vasculature, we next profiled fetal serum samples for circulating growth factors, cytokines, and chemokines to define humoral cues altered in association with impaired placental function. We observed a large (∼2.5-fold) increase in circulating fetal prolactin levels with HFHS diet in both male and female fetuses. Circulating IL-6 levels were also elevated in HFHS fetuses, with a significant male bias (main effect of sex, p = 0.023); while the difference in females was pronounced, only the difference between CON and HFHS males reached statistical significance (p = 0.025; <u>Figure 7a</u>). Levels of chemokines CCL2 and CXCL1 were also moderately elevated in HFHS compared to CON fetuses (main effect of diet, p = 0.012 and 0.045, respectively), with a significant posthoc difference in males only for CCL2 (p = 0.048). Levels of circulating leptin, PlGF, TNF, EGF, HGF, FGF-2, and soluble endoglin (sENG), were not significantly different between groups (<u>Figure 7a</u>). Interestingly, fetal serum prolactin, IL-6, CCL2, and CXCL1 levels were all strongly correlated with one another in HFHS samples, suggesting a shared role in fetal response to maternal HFHS diet and associated placental inflammation (<u>Supplementary Figure 3</u>).

**Figure 7.**
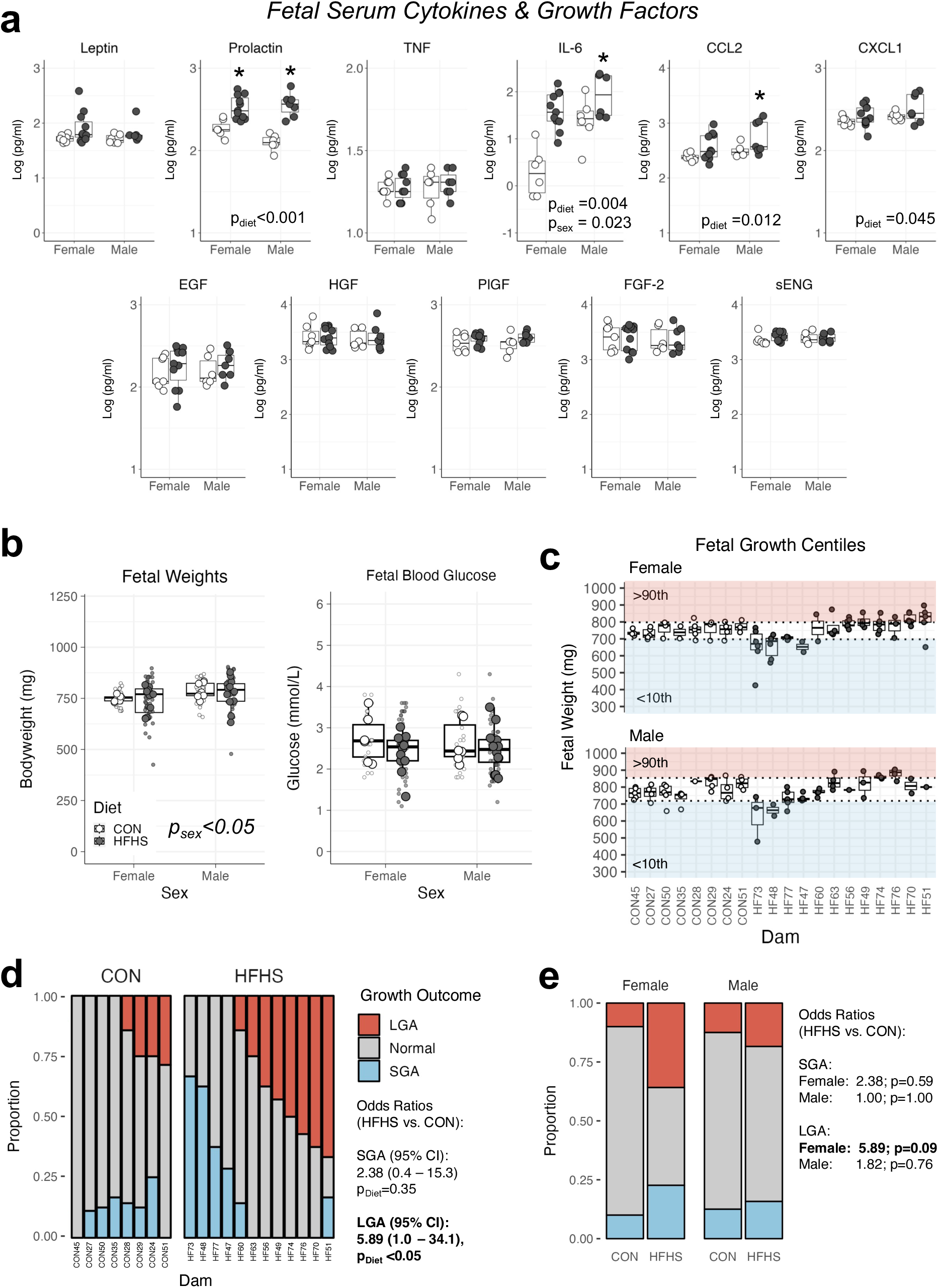
Impacts of maternal HFHS diet on circulating fetal growth factors and inflammatory mediators. **(a)** Multiplex profiling of inflammatory, endocrine, and angiogenic mediators in serum samples pooled by sex from CON (n = 6 – 7/sex) and HFHS (n = 7 – 11/sex) fetuses using magnetic bead immunoassay. **(b)** Fetal bodyweights (left; CON n = 30 – 32/sex, HFHS n = 38 – 54/sex) and blood glucose (right; CON n = 19 – 21/sex, HFHS n = 36 – 53/sex) grouped by litter (CON n= 6 – 8/measure, HFHS n = 12/measure) and sex. **(c)** Fetal bodyweights stratified by litter (dam) and grouped by sex. Sex-specific 10^th^ and 90^th^ centiles for fetal growth were determined based off CON fetal bodyweights (hashed lines) and used to denote growth outcomes according to small for gestational age (SGA; <10^th^ centile, blue) or large for gestational age (LGA; >90^th^ centile, red). **(d)** Odds of each growth outcome (SGA or LGA) were compared for CON and HFHS using nested multivariable binary logistic regression to calculate Odds Ratios. **(e)** Post-hoc pairwise comparisons of estimated marginal means between CON and HFHS fetuses for SGA and LGA outcomes according to sex. Data in (a – c) are presented with boxplots depicting the IQR (box), median (center line), and min and max values within 1.5 IQR (whiskers). In (b) small data points are representative of a single fetus, large data points depict the average value per litter for fetuses of the same sex. Data were analysed using two-way ANOVA with post-hoc Bonferroni-adjusted pairwise comparisons (a), two-way ANOVA of mixed-effects linear models with diet and sex as fixed effects and dam as a random effect (b), or nested binary logistic regression based on generalized linear mixed effect modelling, using diet and sex as fixed effects and dam as a random effect (d) with post-hoc pairwise comparisons using estimated marginal means (e). P-values are indicated on plots where main effects were statistically significant. * Indicates p<0.05 for pairwise comparisons. TNF = Tumor necrosis factor; IL-6 = Interleukin 6, CCL2 = chemokine C-C motif ligand 2; CXCL1 = chemokine C-X-C motif ligand 1; EGF = Epidermal growth factor; HGF = Hepatocyte growth factor, PlGF = Placental growth factor isoform 2; FGF-2 = Fibroblast growth factor 2; sENG = soluble Endoglin.

### HFHS diet increases the odds of fetal overgrowth in female fetuses

Imbalances in fetoplacental oxygen demands and maternal provision can result in hypoxia, leading to fetal growth restriction and/or long-term health impairments (24, 128), whereas macronutrient excess can lead to increased nutrient storage as fat, and increased birth weight (8, 106). Despite the cumulative evidence of placental malperfusion in this model, we observed no differences in mean fetal body weight or blood glucose between diet groups (<u>Figure 7b</u>). However, we observed a statistically significant increase in the variance of fetal body weights within HFHS pregnancies (F_90,61_= 3.11, p < 0.001). In our analysis, we accounted for intra-litter variance in fetal weight (129) and observed significant variability in fetal weight responses to HFHS diet. We categorized fetuses as “small for gestational age” (SGA) or “large for gestation age” (LGA), according to whether they fell below or above the 10^th^ or 90^th^ centiles of CON fetal weights, respectively (39, 95). Centiles were calculated and applied to each sex separately, because of the difference in fetal weights between males and females (39, 130) (main effect of sex, p = 0.001; <u>Figure 7b</u>). When examining these outcomes according to litter, a clear intra-litter similarity in fetal growth outcomes emerged, with some HFHS pregnancies being more prone to SGA and others LGA (<u>Figure 7c</u>). Using nested binary logistic regression to account for repeated measurements within each pregnancy, we found significantly higher odds of LGA fetuses in HFHS pregnancies compared to CON (Odds-ratio: 5.9 (95% CI 1.0 – 34.1), p = 0.048, <u>Figure 7d, Supplementary Table 5</u>). The odds of SGA in contrast, were not significantly different between CON and HFHS pregnancies (Supplementary Table 6). Pairwise comparisons revealed that the effect of HFHS on LGA outcome was driven by female fetuses (Odds ratios: 5.9 (CI: 0.8 – 43.6) vs. 1.8 (CI: 0.2 – 14.7), p = 0.093 and 0.756, for females and males, respectively; <u>Figure 7e</u>). Collectively, these data highlight that the impact of diet-induced excess adiposity on fetal growth varies considerably within and between pregnancies and is modified according to fetal sex.

## DISCUSSION

In this study, we investigated the impacts of a periconceptional obesogenic diet and its associated excess adiposity and dysglycemia on placental oxygenation, vascular adaptations, tissue pathology, and fetal outcomes in late gestation. We show that chronic HFHS feeding results in increased maternal weight, hyperglycemia, and hyperinsulinemia before pregnancy. Once pregnant, HFHS dams exhibited an altered pattern of metabolic, endocrine, and inflammatory adaptations to pregnancy compared to CON-fed females, including reduced gestational weight gain, and reduced levels of stereotypical ‘obesity-associated’ proinflammatory mediators near term. Placentas of HFHS pregnancies were comparatively hypoxic in the labyrinth and junctional zones in late gestation compared to controls. Hypoxia within the labyrinth was coincident with histopathological changes that normally accompany both malperfusion and inflammation, including fibrinoid deposition and calcification, the latter coinciding with focal ablation of fetoplacental vascular membranes and macrophage infiltration. These changes were present in placentas of both sexes, with calcification more strongly associated with HFHS male sex. Consistent with this we observed male-biased overexpression of factors related to fibrosis/calcification in placental labyrinth tissue. Conversely, females demonstrated specific shifts in type 2/regulatory immunity, factors related to tissue repair, and immune cell chemotaxis. This coincided with systemic inflammatory shifts in HFHS fetuses, including elevated levels of soluble inflammatory mediators (IL-6, CCL2, CXCL1, and prolactin). Finally, we observed an overall increased odds of fetal overgrowth in HFHS pregnancies, especially in female fetuses. These fetal and placental outcomes are consistent with changes seen in pregnancies of overweight or obese individuals and primate models of prolonged western diet exposure, where offspring birthweights are similar to or larger than that of lean counterparts, but still exhibit signs of reduced placental efficiency, malperfusion, and inflammation (18, 30, 33–35, 40, 75, 131–133).

Previous studies have suggested that *fetal* hypoxemia is a likely driver of adaptations and adverse outcomes in the context of obesity and gestational diabetes mellitus (GDM) (15, 17, 19, 21, 24, 25). Using an oxygen-sensitive tracer, pimonidazole, we have definitively demonstrated that *placental* oxygen saturation is reduced in pregnancies complicated by maternal HFHS diet in both labyrinth and junctional zone tissue in late gestation. The presence of hypoxia within the placental labyrinth, the first layer perfused by maternal blood in mice, suggests that there is an overall imbalance between maternal oxygen supply and fetoplacental demands in HFHS pregnancies in late gestation. While oxygen supply may be reduced, we did not observe gross reductions to fetal growth, placental ultrastructure, or increases in HIF-1α expression observed in some previous studies reporting signs of placental hypoxia (14–17, 19). This raises new questions as to the severity, chronicity, and etiology of hypoxia in our model.

One such hypothesis is that placental oxygenation is chronically reduced, but not compromised enough to induce angiogenic adaptations, growth restriction, or fetal loss. Chronic reductions in uteroplacental blood flow have been reported in primate models of chronic western (i.e., HFHS) diet feeding without pronounced changes in placental morphometry or fetal growth outcomes (33–35). Moderate fetal hypoxemia has also been observed in human obese pregnancies at term without any impacts on term fetal weight (20). A considerable reserve capacity in placental oxygen and nutrient uptake exists to buffer transient bouts of deprivation and/or loss of villous tissue before fetal growth is compromised (24, 45, 134). Several fetal adaptations, including increases in erythropoietin and hemoglobin levels also occur to enhance oxygen extraction capacity before placental structural adaptations are required to compensate for hypoxia (24). However, this reserve capacity is finite and additional insults beyond those that induce oxygen deprivation can result in frank placental dysfunction and fetal compromise. Thus, our data might suggest that even though this relative hypoxic state has no severe consequences for fetal or placental growth, the capacity to tolerate additional insults could be impaired.

An alternate hypothesis is that this placental hypoxic state is of recent onset and may worsen with advancing gestation. The labyrinth proteome showed increases in IGFBP-1 and -3 in HFHS placentas; these are oxygen-sensitive proteins and act to limit fetal growth by sequestering mitogenic insulin-like growth factors. This could suggest that processes to slow fetal growth are active in HFHS pregnancies. Late-onset placental insufficiency may arise once fetoplacental demands eclipse placental transfer capacity and is thought to underlie complications in late pregnancy to which obese individuals are more predisposed, including term preeclampsia (PE), late-onset fetal growth restriction (FGR) or stillbirth (4, 35, 135). We observed reduced circulating placental growth factor levels (PlGF) in pregnant HFHS dams. Placental growth factor has previously been used in combination with other circulating biomarkers like sFLT-1 (i.e., elevated sFLT-1: PlGF) to non-invasively predict placental dysfunction before the onset of disorders like PE and fetal growth restriction (136, 137). While we were unable to assess circulating sFLT-1 levels, we hypothesize that reduced circulating PlGF is evidence of declining placental function in HFHS pregnancies.

Placental hypoxia in HFHS pregnancies coincided with fibrinoid-type depositions, and placental calcifications overlapping with regions of dystrophic tissue damage. Similar patterns of placental calcification have been seen in other murine models of diet-induced excess adiposity (39, 64) and both increased placental calcifications and deposition of fibrin within the maternal blood spaces of the placenta have been reported in primate models of chronic western diet (i.e., HFHS) feeding (33–35). In these models, histopathologic lesions of calcification and perivillous fibrin deposition co-occurred with increases in uterine artery pulsatility, resistance, and slowed transit of maternal blood through the placenta (35). Reductions in uteroplacental blood flow can lead to increased coagulation, stasis and/or infarction of maternal blood flow through the placenta, contributing to reduced placental oxygen extraction and transfer, or focal ischemia and tissue necrosis (123). This is consistent with our findings of focal loss of both syncytiotrophoblast and fetoplacental vascular tissue in regions of calcification. This leads us to hypothesize that impairments in uteroplacental blood flow could be an upstream driver of placental hypoxia and the pathological lesions in our model. Uteroplacental perfusion is influenced by several factors, including maternal systemic cardiovascular adaptations to pregnancy, uterine vascular remodelling of spiral arteries, and local placental inflammatory processes (42). Future studies of maternal and uteroplacental hemodynamics throughout pregnancy, along with histological assessments of spiral artery remodelling are needed to evaluate their contributions to placental dysfunction in HFHS pregnancies.

While we cannot currently discern exactly how each of these pathological changes to the placenta arise in our model, we suspect that some may arise independently of maternal malperfusion. For example, recent work by Hufnagel et al. demonstrated that the burden of placental calcification in diet-induced obese mice is unaffected by treatments which improve uteroplacental hemodynamics in late pregnancy (64). We suspect that local placental inflammation could instead be driving these changes. We found that HFHS placentas exhibit increases in expression of acute phase proteins (SAP, CRP) which opsonize dying cells and extracellular pathogens, facilitating complement activation and clearance by macrophages (138). We have shown for the first time that regions of calcification in HFHS placentas coincide with focal accumulation of macrophages. Macrophage-dense foci are seen in cases of chronic inflammation of placental villi (chronic villitis/villitis of unknown etiology (VUE; (139, 140) and the chorionic plate and membranes (chorioamnionitis) in human pregnancy (141). These pathologies occur more frequently in pregnancies complicated by maternal obesity (18, 40, 133, 142, 143). Moreover, inflammatory processes associated with VUE involve both maternal T-cell infiltration and destruction of fetoplacental vasculature. We found that calcified regions had residual GS-Iβ4 lectin reactivity for vascular basement membranes with an absence of staining for syncytiotrophoblast and endothelial cells. While we have yet to probe for the presence of T-cells, several T-cell-associated chemokines and cytokines were altered in HFHS labyrinth tissue, suggesting possible similarities to the inflammatory pathogenesis of VUE seen in humans.

There are several possible triggers for labyrinth tissue inflammation in HFHS pregnancies. In both human and animal models, obesity has been linked to heightened placental oxidative stress (39, 144–147) and impaired autophagy (148, 149) leading to cellular damage, apoptosis, and inflammation, which are likely to trigger the recruitment of macrophages (150). Maternal obesity has also been associated with a state of systemic low-grade endotoxemia, including during pregnancy (28). While we do not know whether HFHS dams exhibit increased circulating LPS levels we have previously shown in other work that maternal high-fat feeding does increase circulating LPS levels at term (15). Moreover, recent work in a preconception HFHS feeding model like ours shows that placentas of HFHS pregnancies contain increased levels of lipopolysaccharide (LPS) (151). Pregnant mice administered LPS exhibit placental fibrinoid necroses and mineralization, and have higher placental expression of *Il6* and *Cxcl1*, mirroring both the placental pathology and fetal cytokine profiles seen in our HFHS model (127). Moreover, our data are consistent with previous reports that increases in fetal IL-6 levels are associated with HFHS feeding and are accompanied by elevated IL-17, which directs immune responses against extracellular pathogens (152). We extend these findings to show that HFHS fetuses have increased levels of prolactin, which is stimulated by both LPS and IL-6 (153, 154) and has been shown to attenuate TLR4-driven inflammation in the placenta and fetal membranes (155, 156). Furthermore, prolactin has been previously shown to be elevated in human fetuses from pregnancies complicated by pregnancy-induced hypertension (i.e., PE) (157). LPS administration (albeit at higher doses) is a common preclinical model of PE, stimulating systemic coagulation which impairs uteroplacental blood flow (158–160). It is possible that a spectrum of similar inflammatory processes could underlie both inflammatory and hypoxic changes we observe in HFHS placentas. Our findings also reinforce that fetoplacental outcomes in HFHS pregnancies vary depending on sex. Labyrinth expression of several cytokines and chemokines related to inflammation (CRP), tissue repair (IL-33, IL-4), and T-cell recruitment/cytokine production (CCL17, IL-27) were increased in female but not male placentas. This data suggests that HFHS female fetuses exhibit an overall greater immune response to stressors associated with maternal HFHS diet. This aligns with recent human data showing that female placental macrophages exhibit a more pronounced response to inflammatory stimuli (161) and that female fetuses generally invest more in maintaining placental homeostasis while males prioritize fetal growth at the risk of greater morbidity and mortality (162). Previous work in HFHS pregnancies has shown that a heightened state of placental inflammation in males is linked to sex-specific consequences for offspring behavioural phenotypes (151). In our study, females showed a lesser degree of placental calcification, and a greater degree of fetal overgrowth compared to males. Sex-specific adaptations driven by placental immune cells in response to the challenges posed by maternal HFHS diet may contribute to divergent outcomes for fetal growth and offspring health.

The humoral cues propagated to fetuses likely have lasting consequences for offspring health. Elevations in fetal IL-6 alone are sufficient to program long-term impairments in offspring, with T_h_17 skewing in intestinal immune profiles that might predispose to gut dysbiosis and inflammation (163). We have also previously shown that fetuses of mice fed a very high-fat diet (60% kCal from fat) have increased NFκB activity and alterations to gut barrier proteins at term gestation (15), and others have demonstrated increased T_h_17 inflammation in high-fat diet-fed offspring challenged with DSS-induced colitis (164). Elevated fetal IL-6 levels, including in the context of diet-induced obesity, have also been shown to impair the formation of neural circuits that regulate offspring feeding and social behaviours (165–167), potentially linking fetoplacental inflammation to changes in postnatal appetite and cognition in offspring of pregnancies complicated by maternal obesity in humans (13), as well as offspring of HFHS diet-fed dams (151). We recognize that our study is not without limitations. Pimonidazole immunostaining represents one of the most accessible and accurate ways of comparing tissue oxygenation between experimental groups. However, these data are cross-sectional and do not provide a direct measure of oxygen saturation (i.e., pO_2_) or its kinetics over time. While reduced labyrinth oxygenation did not coincide with the global upregulation of HIF-1α protein levels, our measures do not account for cellular localization and are relative to total protein. Other HIF family membranes, including HIF-2α, are expressed within the placental labyrinth and may mediate cellular responses to more moderate hypoxia (47, 168, 169) and reduced uteroplacental perfusion (170). The disparity in oxygen saturation between placental zones could also indicate that HIFs are solely activated in the junctional zone. Junctional zone-specific activation of HIF-1α has been shown to produce features of preeclampsia in mice (169), which may have relevance for our findings in this study but has yet to be investigated. Immunohistochemical localization of both HIF-1α and HIF-2α within placental tissues could ascertain the regulation of placental HIF signalling in metabolically complicated pregnancies. Our analyses of placental morphology and vascular structure do not capture other physiologically relevant changes to the placenta that can occur in the context of hypoxia (39, 171), including changes to blood vessel branching, interhaemal barrier thickness, and organization of maternal blood spaces. Moreover, while total vascular areas were similar between groups, we did not account for areas affected by calcification or fibrin deposition, which could impair overall capacity for diffusion. Stereological evaluation of these measures and volumetric changes to the placenta is needed to account for this, as has been done in other models (39).

In conclusion, we show that maternal HFHS diet-induced excess adiposity is associated with placental hypoxia and coincides with histological patterns of malperfusion and inflammation. These placental impairments contribute to changes in pathways that regulate fetal growth that likely have impacts on offspring health. While we see considerable evidence of maternal malperfusion-related injury to the placenta in HFHS pregnancies, and given the known risk of hypertensive disorders of pregnancy in pregnancies complicated by obesity (6), future work should include additional measures of maternal perfusion of the placenta as well as profiling of maternal blood pressure, indices of uterine artery function, and uterine spiral artery remodelling across gestation.

## AUTHOR CONTRIBUTIONS

CJB and DMS conceived the study. DMS acquired funding. CJB, TAR, EY, PAJ, HL, and JB collected data. TAR, EY, PAJ, HL, and JB provided technical assistance. CJB performed all formal analyses and data visualization. CJB and DMS interpreted the data. CJB drafted the manuscript. CJB and DMS edited the manuscript. DMS provided resources and supervision for this study. All authors have read and approved the manuscript in its final form.

## ACKNOWLEDGEMENTS

The authors would like to thank Dr. Sue Ozanne and Dr. Antonia Hufnagel for their kind provision of the tissue calcium staining protocol. CJB was supported by a Fredrick Banting and Charles Best Doctoral Award from the Canadian Institutes of Health Research. DMS is supported by the Canada Research Chairs Program. This work was supported by a Project Grant awarded to DMS by the Canadian Institutes of Health Research.

