## Supplemental Tables and Figures for "Maternal high-fat, high-sucrose diet-induced excess adiposity is linked to placental hypoxia and disruption of fetoplacental immune homeostasis in late gestation"

Bellissimo, CJ *et al.* (2024b)

### SUPPLEMENTARY MATERIALS

**Supplementary Table 1**. Nutrient composition comparison of experimental diets

<table>
<tr><th>Nutritional Value</th><th>CON – Teklad 22/5 Rodent Diet</th><th>HFHS – Research Diets D12451</th></tr>
<tr><td>Energy Density (kCal/g)</td><td>3.00</td><td>4.73</td></tr>
<tr><td colspan="3">Macronutrients – % kCal/g</td></tr>
<tr><td>Fat[1,2]</td><td>17%</td><td>45%</td></tr>
<tr><td>Sucrose</td><td>NA</td><td>17%</td></tr>
<tr><td>Carbohydrate[3]</td><td>54%</td><td>18%</td></tr>
<tr><td>Protein[4]</td><td>29%</td><td>20%</td></tr>
<tr><td colspan="3">Supplemented Vitamins – sources[5]</td></tr>
<tr><td>Vitamin A</td><td>Vitamin A</td><td>Vitamin A acetate</td></tr>
<tr><td>Vitamin $B_1$</td><td rowspan="9">Dietary[5]</td><td>Thiamine HCl</td></tr>
<tr><td>Vitamin $B_2$</td><td>Riboflavin</td></tr>
<tr><td>Vitamin $B_3$</td><td>Nicotinic acid</td></tr>
<tr><td>Vitamin $B_5$</td><td>Pantothenic acid, d, Calcium</td></tr>
<tr><td>Vitamin $B_6$</td><td>Pyridoxine HCl</td></tr>
<tr><td>Vitamin $B_7$</td><td>Biotin, 1%</td></tr>
<tr><td>Vitamin $B_9$</td><td>Folic acid</td></tr>
<tr><td>Vitamin $B_{12}$</td><td>Cyanocobalamin, 0.1%</td></tr>
<tr><td>Vitamin D</td><td>Vitamin $D_3$</td><td>Vitamin $D_3$</td></tr>
<tr><td>Vitamin E</td><td rowspan="3">Dietary[5]</td><td>Vitamin E Acetate</td></tr>
<tr><td>Vitamin K</td><td>Menadione sodium bisulfite</td></tr>
<tr><td>Choline</td><td>Choline bitartrate</td></tr>
<tr><td colspan="3">Supplemented minerals – sources[6]</td></tr>
<tr><td>Calcium</td><td rowspan="12">Dietary[6]</td><td>Calcium carbonate</td></tr>
<tr><td>Phosphorus</td><td>Calcium phosphate</td></tr>
<tr><td>Sodium</td><td>Sodium chloride</td></tr>
<tr><td>Potassium</td><td>Potassium citrate,</td></tr>
<tr><td>Chloride</td><td>Sodium chloride</td></tr>
<tr><td>Magnesium</td><td>Magnesium sulfate, Magnesium oxide</td></tr>
<tr><td>Copper</td><td>Copper carbonate</td></tr>
<tr><td>Iodine</td><td>Potassium iodate</td></tr>
<tr><td>Iron</td><td>Ferric citrate</td></tr>
<tr><td>Manganese</td><td>Manganese carbonate</td></tr>
<tr><td>Selenium</td><td>Sodium selenite</td></tr>
<tr><td>Zinc</td><td>Zinc carbonate</td></tr>
<tr><td>Molybdenum</td><td>Not described</td><td>Ammonium molybdate – $4H_2O$</td></tr>
</table>

[1] Sources – CON: multiple; HFHS: lard, soybean oil (https://insights.envigo.com/hubfs/resources/data-sheets/8640-datasheet-0915.pdf)
[2] Cholesterol content – CON: 30mg/kg; HFHS: 195.5mg/kg (combined lard + casein)
[3] Sources – CON: multiple; HFHS: corn starch, maltodextrin 10
[4] Sources – CON: multiple; HFHS: Casein (mesh 30), L-Cystine
[5] Sources – CON: dietary; HFHS: vitamin mix V10001C (see https://www.researchdiets.com/formulas/V10001C)
[6] Sources – CON: multiple, see Appendix A.1; HFHS: mineral mix S10026B (see https://www.researchdiets.com/formulas/S10026B)

**Supplementary Table 2.** Genotyping primers for embryonic sex determination

| Gene | Amplicon size (bp) | Primer | Sequence (5'-3') |
|---|---|---|---|
| *Sry* | 273 | *Forward* | TTGTCTAGAGAGCATGGAGGGCCATGTCAA |
| | | *Reverse* | CCACTCCTCTGTGACACTTTAGCCCTCCGA |
| *Fabp2* | 194 | *Forward* | TGGACAGGACTGGACCTCTGCTTTCCTAGA |
| | | *Reverse* | TAGAGCTTTGCCACATCACAGGTCATTCAG |

**Supplementary Table 3.** Immunostaining conditions and reagents

| Experiment | Antigen Retrieval | Primary Antibody (Dilution)<br>Vendor- cat. No.,<br>RRID | Isotype Control (Dilution)<br>Vendor- cat. No.,<br>RRID | Secondary Antibody (Dilution)<br>Vendor- cat. No.,<br>RRID |
|---|---|---|---|---|
| Placental hypoxia | Tris-EDTA<br>pH 9.0<br>(95$^{o}$C, 30 min) | Chicken anti-SLC16A1 (1:200)<br>Millipore Sigma AB1286-I<br>AB_90565<br><br>Rabbit anti-Pimonidazole (1:100)<br>Hypoxyprobe – Pab27 | Chicken IgY (1:200)<br>BL 402101<br><br>Rabbit IgG (1:500)<br>TFS – 02-6102,<br>AB_2532938 | Goat anti-Chicken IgY–AF488 (1:500)<br>TFS – A-11039<br>AB_2534096<br><br>Goat anti-Rabbit IgG–AF647 (1:500)<br>TFS – A32733<br>AB_2633282 |
| Placental Vessels | Tris-EDTA<br>pH 9.0<br>(95$^{o}$C, 30 min) | Biotinylated Isolectin $\beta$4 from *Griffonia Simplicifolia* (1:200)<br>VL B1205<br>AB_2314661<br><br>Mouse anti- $\alpha$-SMA (1:250)<br>Abcam – ab7187<br>AB_262054<br><br>Rabbit anti-CD31 (1:50)<br>Abcam – ab28364<br>AB_726362 | NA (Lectin omitted)<br><br>Mouse IgG2a (1:250)<br>TFS – 02-6200<br>AB_2532943<br><br>Rabbit IgG (1:1250)<br>TFS – 02-6102,<br>AB_2532938 | Streptavidin Dylight488 (1:500)<br>VL – SA5488<br><br>Goat anti-Mouse IgG – AF555 (1:500)<br>TFS – A32727<br>AB_2633276<br><br>Goat anti-Rabbit IgG–AF647 (1:500)<br>TFS – A32733<br>AB_2633282 |
| Placental macrophages | Tris-EDTA<br>pH 9.0<br>(95$^{o}$C, 30 min) | Chicken anti-SLC16A1 (1:200)<br>Millipore Sigma AB1286-I<br>AB_90565<br><br>Rabbit anti-F4/80 (1:200)<br>CST – 70076,<br>AB_2799771 | Chicken IgY (1:200)<br>BL 402101<br><br>Rabbit IgG (1:1000)<br>TFS – 02-6102,<br>AB_2532938 | Goat anti-Chicken IgY–AF488 (1:500)<br>TFS – A-11039<br>AB_2534096<br><br>Goat anti-Rabbit IgG – biotin (1:200)<br>VL – VECTPK6101<br><br>Streptavidin AF647 (1:500)<br>TFS – S32357 |

CST = Cell Signalling Technologies
TFS = ThermoFisher Scientific
VL = Vector Laboratories
BL = Biolegend

**Supplemental Table 4.** Mouse Proteome Profiler array of E17.5 placental labyrinth tissue

| Coordinates | Analyte | Protein Name | Relevant aliases | Exposure (seconds) | Background Corrected Volume (Densitometric Units) | | | |
|---|---|---|---|---|---|---|---|---|
| | | | | | *CON Female* | *HFHS Female* | *CON Male* | *HFHS Male* |
| A1/2 | positive control | - | - | - | | | | |
| A3/4 | AdipoQ | Adiponectin | Acrp30 | 480 | 219006 | 252186 | 219694 | 308698 |
| | | | | | 207342 | 266902 | 164842 | 287514 |
| A5/6 | AREG | Amphiregulin | - | 480 | 30706 | 27858 | 36526 | 40114 |
| | | | | | 36674 | 29526 | 37946 | 35714 |
| A7/8 | ANGPT1 | Angiopoetin-1 | ANG1 | 480 | 63982 | 55114 | 71058 | 58842 |
| | | | | | 59634 | 50978 | 61414 | 57886 |
| A9/10 | ANGPT2 | Angiopoetin-2 | ANG2 | 480 | 240330 | 229666 | 228026 | 227258 |
| | | | | | 226794 | 233602 | 219226 | 223406 |
| A11/12 | ANGPTL3 | Angiopoietin-like 3 | ANG5 | 480 | 147398 | 146998 | 189710 | 128438 |
| | | | | | 151006 | 139454 | 188858 | 115642 |
| A13/14 | BAFF | B-cell Activating Factor | TNFSF13B | 480 | 23158 | 19962 | 29394 | 20786 |
| | | | | | 32450 | 29258 | 28954 | 26982 |
| A15/16 | CD93 | Cluster of differentiation 93 | C1QR1 | 120 | 275106 | 240680 | 330422 | 221674 |
| | | | | | 259874 | 243560 | 316962 | 235426 |
| A17/18 | CCL2 | chemokine C-C motif ligand 2 | MCP-1 | 480 | 22166 | 29162 | 29682 | 16458 |
| | | | | | 23506 | 26166 | 27890 | 19170 |
| A19/20 | CCL3/4 | chemokine C-C motif ligand 3/4 | MIP-1 alpha/beta | 480 | 23070 | 26150 | 33398 | 22538 |
| | | | | | 22674 | 26362 | 43158 | 22274 |
| A21/22 | CCL5 | chemokine C-C motif ligand 5 | RANTES | 480 | 21294 | 8666 | 15462 | 27278 |
| | | | | | 27438 | 19042 | 21382 | 44786 |
| A23/24 | positive control | - | - | - | | | | |
| B1/2 | - | - | - | 480 | | | | |

| | | | | | | | | |
|---|---|---|---|---|---|---|---|---|
| B3/4 | CCL6 | chemokine C-C motif ligand 6 | - | 480 | 100426 | 111230 | 96494 | 101862 |
| | | | | | 100090 | 110846 | 95062 | 96626 |
| B5/6 | CCL11 | chemokine C-C motif ligand 11 | Eotaxin-1 | 480 | 74694 | 68782 | 66922 | 74710 |
| | | | | | 77042 | 64142 | 67286 | 74030 |
| B7/8 | CCL12 | chemokine C-C motif ligand 12 | MCP-5 | 480 | 55954 | 59234 | 55922 | 58570 |
| | | | | | 47926 | 53042 | 76638 | 52002 |
| B9/10 | CCL17 | chemokine C-C motif ligand 17 | TARC | 480 | 51118 | 68166 | 80978 | 50838 |
| | | | | | 50510 | 62618 | 88602 | 48570 |
| B11/12 | CCL19 | chemokine C-C motif ligand 19 | MIP-3 beta | 480 | 43626 | 49110 | 69518 | 37854 |
| | | | | | 40794 | 45322 | 71286 | 37682 |
| B13/14 | CCL20 | chemokine C-C motif ligand 20 | MIP-3 alpha | 480 | 17310 | 16886 | 26650 | 9202 |
| | | | | | 24878 | 18446 | 25930 | 14490 |
| B15/16 | CCL21 | chemokine C-C motif ligand 21 | 6CKine | 480 | 180770 | 233366 | 207618 | 260993 |
| | | | | | 186358 | 228958 | 196718 | 218646 |
| B17/18 | CCL22 | chemokine C-C motif ligand 22 | MDC | 480 | 99734 | 78742 | 101698 | 72566 |
| | | | | | 93934 | 77634 | 97022 | 69946 |
| B19/20 | CD14 | Cluster of differentiation 14 | - | 480 | 57658 | 55054 | 63818 | 50246 |
| | | | | | 52202 | 56374 | 64438 | 47574 |
| B21/22 | CD40 | Cluster of differentiation 40 | TNFRSF5 | 480 | 291554 | 262834 | 253622 | 257414 |
| | | | | | 304470 | 276866 | 257510 | 264858 |
| B23/24 | - | - | - | - | | | | |
| C1/2 | - | - | - | - | | | | |
| C3/4 | CD160 | Cluster of differentiation 160 | - | 480 | 35726 | 39686 | 42738 | 42590 |
| | | | | | 34182 | 39838 | 39434 | 40178 |
| C5/6 | RARRES2 | Retinoic acid responder protein 2 | Chemerin, TIG2 | 120 | 138054 | 141588 | 135114 | 155158 |
| | | | | | 146562 | 140700 | 124026 | 147554 |
| C7/8 | CHi3L1 | Chitinase 3-like 1 | - | 480 | 127682 | 125350 | 113602 | 122362 |

| | | | | | | | | |
|---|---|---|---|---|---|---|---|---|
| | | | | | 124750 | 117874 | 121898 | 119086 |
| C9/10 | F3 | Tissue Factor | TF, CD142 | 120 | 437734 | 487892 | 498294 | 421150 |
| | | | | | 401498 | 462332 | 515614 | 417982 |
| C11/12 | C5a | Complement component 5a | C5 | 480 | 45330 | 79034 | 77314 | 67178 |
| | | | | | 31646 | 46926 | 51754 | 36302 |
| C13/14 | CFD | Complement factor D | Adipsin | 480 | 81498 | 70246 | 104262 | 49282 |
| | | | | | 79726 | 75178 | 102914 | 63506 |
| C15/16 | CRP | C-reactive protein | - | 480 | 307850 | 437618 | 369286 | 377046 |
| | | | | | 324122 | 428294 | 359826 | 393538 |
| C17/18 | CX3CL1 | chemokine C-X3-3-motif ligand 1 | Fracktalkine | 480 | 322278 | 317466 | 396366 | 335182 |
| | | | | | 347018 | 319014 | 400110 | 332194 |
| C19/20 | CXCL1 | chemokine C-X-C motif ligand 1 | KC, GRO-alpha | 480 | 24346 | 15766 | 21434 | 15646 |
| | | | | | 8978 | 11630 | 8550 | 11590 |
| C21/22 | CXCL2 | chemokine C-X-C motif ligand 2 | GRO-beta, MIP-2 alpha | 480 | 27650 | 41134 | 49398 | 44250 |
| | | | | | 34530 | 36638 | 52874 | 48174 |
| C23/24 | - | - | - | - | | | | |
| D1/2 | CXCL9 | chemokine C-X-C motif ligand 9 | MIG | 480 | 22322 | 33858 | 28914 | 32110 |
| | | | | | 24626 | 29822 | 34486 | 32090 |
| D3/4 | CXCL10 | chemokine C-X-C motif ligand 10 | IP-10 | 480 | 38118 | 41026 | 49310 | 51226 |
| | | | | | 36634 | 48466 | 49330 | 50922 |
| D5/6 | CXCL11 | chemokine C-X-C motif ligand 11 | I-TAC | 480 | 25298 | 44470 | 31194 | 28162 |
| | | | | | 28018 | 39178 | 35530 | 30566 |
| D7/8 | CXCL13 | chemokine C-X-C motif ligand 13 | BLC, BCA-1 | 480 | 27574 | 39686 | 34138 | 29498 |
| | | | | | 27678 | 37558 | 34058 | 25158 |
| D9/10 | CXCL16 | chemokine C-X-C motif ligand 16 | - | 480 | 75594 | 97514 | 78586 | 79766 |
| | | | | | 76774 | 108282 | 92314 | 94390 |
| D11/12 | CST3 | Cystatin 3 | - | 120 | 170358 | 177892 | 209238 | 153950 |
| | | | | | 158650 | 164180 | 190674 | 136978 |
| D13/14 | DKK-1 | | - | 480 | 65710 | 63118 | 65730 | 53370 |

| | | | | | | | | |
|---|---|---|---|---|---|---|---|---|
| | | Dickkopf-related protein 1 | | | 67366 | 59970 | 71150 | 51346 |
| D15/16 | DPP4 | Dipeptidyl peptidase 4 | DPPIV, CD26 | 480 | 105070 | 90162 | 88710 | 75550 |
| | | | | | 93658 | 92546 | 98102 | 49294 |
| D17/18 | EGF | Epidermal growth factor | - | 480 | 26354 | 33634 | 36658 | 32186 |
| | | | | | 31038 | 37742 | 40386 | 25730 |
| D19/20 | Endoglin | - | ENG, CD105 | 120 | 183126 | 188708 | 177302 | 136638 |
| | | | | | 182810 | 191160 | 215798 | 138510 |
| D21/22 | Endostatin | - | - | 120 | 204162 | 254772 | 290222 | 270238 |
| | | | | | 213934 | 248220 | 307018 | 267874 |
| D23/24 | AHSG | Alpha-2-HS-glycoprotein | Fetuin A | 120 | 206478 | 225712 | 270850 | 251574 |
| | | | | | 199970 | 242596 | 302586 | 240762 |
| E1/2 | FGF-1 | Fibroblast growth factor 1 | aFGF | 480 | 61990 | 71130 | 40170 | 91978 |
| | | | | | 63970 | 70650 | 75906 | 89262 |
| E3/4 | FGF-21 | Fibroblast growth factor 21 | - | 480 | 28502 | 33742 | 40966 | 38566 |
| | | | | | 24274 | 32710 | 35238 | 36018 |
| E5/6 | FLT3LG | FLT-3 Ligand | FLT3L | 480 | 35434 | 34702 | 42678 | 34918 |
| | | | | | 41250 | 35594 | 45222 | 31462 |
| E7/8 | GAS6 | Growth arrest-specific 6 | - | 480 | 42182 | 51550 | 56442 | 39542 |
| | | | | | 40370 | 51034 | 54118 | 38578 |
| E9/10 | CSF3 | Granulocyte colony stimulating Factor | G-CSF | 480 | 38906 | 38574 | 51470 | 38410 |
| | | | | | 35482 | 34270 | 45490 | 31590 |
| E11/12 | GDF15 | Growth/differentiation factor 15 | - | 480 | 30334 | 41246 | 37070 | 25742 |
| | | | | | 26130 | 37998 | 35438 | 22118 |
| E13/14 | CSF2 | Granulocyte macrophage colony stimulating factor | GM-CSF | 480 | 9686 | 11614 | 10438 | 7154 |
| | | | | | 8270 | 11914 | 8722 | 5870 |
| E15/16 | HGF | Hepatocyte growth factor | - | 480 | 45766 | 43530 | 47554 | 31254 |
| | | | | | 45922 | 43578 | 43654 | 32318 |
| E17/18 | ICAM-1 | Intercellular adhesion molecule 1 | CD54 | 480 | 421210 | 478881 | 411002 | 301790 |
| | | | | | 434926 | 493165 | 418030 | 311614 |

| | | | | | | | | |
|---|---|---|---|---|---|---|---|---|
| E19/20 | IFN-γ | Interferon gamma | - | 480 | 34398 | 43994 | 38038 | 27774 |
| | | | | | 29794 | 38158 | 33946 | 25374 |
| E21/22 | IGFBP-1 | IGF binding protein 1 | - | 480 | 217990 | 474978 | 315770 | 417698 |
| | | | | | 232746 | 487414 | 352634 | 546090 |
| E23/24 | IGFBP-2 | IGF binding protein 2 | - | 120 | 382902 | 459476 | 578066 | 532526 |
| | | | | | 407462 | 527584 | 626728 | 538966 |
| F1/2 | IGFBP-3 | IGF binding protein 3 | - | 480 | 66082 | 104290 | 74146 | 107710 |
| | | | | | 69446 | 98602 | 88478 | 113666 |
| F3/4 | IGFBP-5 | IGF binding protein 5 | - | 480 | 37738 | 44066 | 46482 | 56478 |
| | | | | | 36546 | 42874 | 45986 | 55182 |
| F5/6 | IGFBP-6 | IGF binding protein 6 | - | 120 | 154134 | 169948 | 159070 | 169510 |
| | | | | | 168686 | 162736 | 165778 | 168530 |
| F7/8 | IL-1a | Interleukin 1 alpha | - | 480 | 68506 | 77654 | 81626 | 67378 |
| | | | | | 59106 | 73438 | 76010 | 61506 |
| F9/10 | IL-1b | Interleukin 1 beta | - | 480 | 24234 | 19046 | 29174 | 19726 |
| | | | | | 22742 | 18978 | 30618 | 20070 |
| F11/12 | IL-1RA | Interleukin 1 receptor agonist | IL1RN | 480 | 49226 | 50230 | 53830 | 39918 |
| | | | | | 43166 | 50174 | 49390 | 37158 |
| F13/14 | IL-2 | Interleukin 2 | - | 480 | 1630 | 1194 | 2858 | 414 |
| | | | | | 1994 | 1858 | 3066 | 1214 |
| F15/16 | IL-3 | Interleukin 3 | - | 480 | 7322 | 4330 | 4162 | 2902 |
| | | | | | 5610 | 5046 | 8426 | 4994 |
| F17/18 | IL-4 | Interleukin 4 | - | 480 | 47314 | 62622 | 53574 | 41342 |
| | | | | | 47014 | 65686 | 51462 | 43250 |
| F19/20 | IL-5 | Interleukin 5 | - | 480 | 23974 | 30066 | 31610 | 20390 |
| | | | | | 23866 | 25398 | 32286 | 22390 |
| F21/22 | IL-6 | Interleukin 6 | - | 480 | 9262 | 9290 | 14698 | 10394 |
| | | | | | 7918 | 9826 | 19470 | 13294 |
| F23/24 | IL-7 | Interleukin 7 | - | 480 | 50018 | 58650 | 79018 | 69090 |
| | | | | | 39670 | 61742 | 80390 | 69158 |

| | | | | | | | | |
|---|---|---|---|---|---|---|---|---|
| G1/2 | IL-10 | Interleukin 10 | - | 480 | 33458 | 42586 | 41026 | 37578 |
| | | | | | 34986 | 41670 | 46830 | 40418 |
| G3/4 | IL-11 | Interleukin 11 | - | 480 | 32182 | 39250 | 40966 | 43794 |
| | | | | | 32786 | 44046 | 47918 | 43370 |
| G5/6 | IL-12 p40 | Interleukin 12 p40 subunit | IL12B | 480 | 45202 | 46154 | 44466 | 45586 |
| | | | | | 42454 | 45618 | 62406 | 43842 |
| G7/8 | IL-13 | Interleukin 13 | - | 480 | 33262 | 47090 | 53934 | 32806 |
| | | | | | 32534 | 37286 | 45506 | 29282 |
| G9/10 | IL-15 | Interleukin 15 | - | 480 | 80350 | 72010 | 87162 | 77390 |
| | | | | | 77126 | 69742 | 86990 | 88910 |
| G11/12 | IL-17a | Interleukin 17a | - | 480 | 3586 | 2470 | 4810 | 1218 |
| | | | | | 1902 | 814 | 2862 | 3270 |
| G13/14 | IL-22 | Interleukin 22 | - | 480 | 17166 | 19494 | 18530 | 12274 |
| | | | | | 15898 | 16518 | 18002 | 11266 |
| G15/16 | IL-23 | Interleukin 23 | - | 480 | 23334 | 22518 | 25838 | 15054 |
| | | | | | 21738 | 21414 | 23422 | 14094 |
| G17/18 | IL-27 p28 | Interleukin 27 p28 subunit | - | 480 | 62494 | 106690 | 76958 | 49594 |
| | | | | | 66562 | 85778 | 80346 | 48014 |
| G19/20 | IL-28 A/B | Interleukin 28 A/B | IFNL2/3 | 480 | 95854 | 126430 | 112262 | 66742 |
| | | | | | 90038 | 116158 | 112534 | 60834 |
| G21/22 | IL-33 | Interleukin 33 | - | 480 | 108342 | 144930 | 184250 | 160210 |
| | | | | | 121554 | 147674 | 165358 | 158718 |
| G23/24 | LDL-R | LDL Receptor | - | 480 | 162306 | 230018 | 249346 | 269646 |
| | | | | | 151674 | 235734 | 244758 | 248218 |
| H1/2 | Leptin | - | - | 480 | 76490 | 116426 | 88814 | 116350 |
| | | | | | 74690 | 111166 | 88814 | 123338 |
| H3/4 | LIF | Leukemia inhibitory factor | - | 480 | 68638 | 51854 | 81374 | 62642 |
| | | | | | 64446 | 61134 | 76466 | 60638 |
| H5/6 | LCN2 | Lipocalin 2 | NGAL | 480 | 168086 | 130886 | 133026 | 121066 |
| | | | | | 177818 | 135126 | 142110 | 125058 |

| | | | | | | | | |
|---|---|---|---|---|---|---|---|---|
| H7/8 | CXCL5 | chemokine C-X-C motif 5 | ENA78, LIX | 480 | 94410 | 83870 | 81802 | 91466 |
| | | | | | 99250 | 81770 | 86630 | 99434 |
| H9/10 | CSF1 | Macrophage colony stimulating factor | M-CSF | 480 | 79666 | 63670 | 67590 | 65066 |
| | | | | | 78890 | 64030 | 78338 | 65726 |
| H11/12 | MMP-2 | Matrix Metalloproteinase 2 | - | 480 | 180106 | 133610 | 152626 | 147090 |
| | | | | | 179798 | 135986 | 145662 | 138602 |
| H13/14 | MMP-3 | Matrix Metalloproteinase 3 | - | 480 | 16734 | 13434 | 20006 | 16094 |
| | | | | | 17402 | 16546 | 19590 | 18046 |
| H15/16 | MMP-9 | Matrix Metalloproteinase 9 | - | 480 | 117950 | 93158 | 95582 | 99546 |
| | | | | | 130370 | 97162 | 104678 | 101254 |
| H17/18 | MPO | Myeloperoxidase | - | 120 | 193158 | 209264 | 220510 | 197694 |
| | | | | | 194742 | 197380 | 222822 | 199290 |
| H19/20 | SPP-1 | Secreted phosphoprotein 1/ Osteopontin | OPN | 480 | 346322 | 436346 | 254510 | 374490 |
| | | | | | 329898 | 425858 | 261086 | 356026 |
| H21/22 | OPG | Osteoprotegerin | TNFSF11b | 120 | 431914 | 509092 | 198162 | 275754 |
| | | | | | 446462 | 523456 | 199790 | 296570 |
| H23/24 | TYMP | Thymidine Phosphorylase | PD-ECGF | 480 | 36114 | 32798 | 32010 | 40630 |
| | | | | | 22006 | 29690 | 33002 | 34646 |
| I1/2 | PDFG-B | Platelet derived growth factor beta | - | 480 | 61514 | 52834 | 64730 | 58138 |
| | | | | | 64990 | 47254 | 67062 | 59014 |
| I3/4 | SAP | Serum amyloid P component | APCS, PTX-2 | 480 | 142258 | 251530 | 160126 | 244670 |
| | | | | | 137674 | 240590 | 159342 | 259778 |
| I5/6 | PTX-3 | Pentraxin 3 | - | 480 | 50422 | 40466 | 55050 | 42410 |
| | | | | | 56322 | 44618 | 53378 | 47094 |
| I7/8 | POSTN | Periostin | OSF-2 | 480 | 40646 | 22402 | 39586 | 29142 |
| | | | | | 42510 | 24094 | 43158 | 26286 |
| I9/10 | DLK-1 | Delta-like non-canonical Notch ligand 1 | - | 120 | 232430 | 197008 | 193110 | 199394 |
| | | | | | 238598 | 198992 | 187962 | 198978 |
| I11/12 | PRL2C2 | Proliferin | PLF, PLF-1 | 120 | 186846 | 166404 | 152242 | 172122 |

| | | | | | | | | |
|---|---|---|---|---|---|---|---|---|
| | | | | | 172126 | 163640 | 148374 | 159794 |
| I13/14 | PCSK9 | Proprotein convertase subtilisin/kexin type 9 | - | 480 | 178466 | 186118 | 179702 | 224518 |
| | | | | | 193538 | 179038 | 170234 | 233962 |
| I15/16 | RAGE | Receptor for advanced glycation end products | AGER | 120 | 191162 | 183352 | 173110 | 199654 |
| | | | | | 191450 | 173228 | 179078 | 180658 |
| I17/18 | RBP4 | Retinol binding protein 4 | - | 480 | 368158 | 346210 | 331278 | 326298 |
| | | | | | 366210 | 337862 | 338566 | 331914 |
| I19/20 | REG3G | Regenerating islet-derived protein 3 gamma | - | 480 | 244250 | 99058 | 194494 | 112358 |
| | | | | | 231126 | 96354 | 203446 | 105610 |
| I21/22 | Resistin | - | RETN, ADSF, XCP1 | 480 | 83806 | 61374 | 62050 | 68502 |
| | | | | | 84690 | 64634 | 67142 | 71262 |
| I23/24 | - | - | - | - | | | | |
| J1/2 | positive control | - | - | - | | | | |
| J3/4 | E-Selectin | - | CD62E | 480 | 296494 | 358950 | 307814 | 354362 |
| | | | | | 307762 | 354458 | 321982 | 344710 |
| J5/6 | P-selectin | - | CD62P | 480 | 118350 | 111658 | 103966 | 109018 |
| | | | | | 152214 | 128866 | 125822 | 149478 |
| J7/8 | PAI-1 | Plasminogen activator inhibitor 1 | SERPINE1 | 120 | 600568 | 572551 | 462334 | 637586 |
| | | | | | 603744 | 572635 | 459738 | 596864 |
| J9/10 | PEDF | Pigment epithelium-derived growth factor | SERPINF1 | 480 | 64786 | 54902 | 79622 | 59310 |
| | | | | | 65598 | 55070 | 58070 | 54778 |
| J11/12 | THPO | Thrombopoetin | - | 480 | 35674 | 33318 | 35818 | 33714 |
| | | | | | 34294 | 31346 | 32302 | 33358 |
| J13/14 | TIM-1 | T-cell immunoglobuin and mucin domain 1 | HAVCR1, KIM-1 | 480 | 500272 | 398122 | 401470 | 407122 |
| | | | | | 478697 | 373946 | 390954 | 402522 |
| J15/16 | TNF | | TNFa | 480 | 49658 | 40038 | 45478 | 50182 |

| | | | | | | | | |
|---|---|---|---|---|---|---|---|---|
| | | Tumor Necrosis Factor | | | 42366 | 41198 | 47894 | 37758 |
| J17/18 | VCAM-1 | Vascular cell adhesion protein 1 | CD106 | 480 | 183918 | 157550 | 186554 | 168678 |
| | | | | | 181810 | 151518 | 183994 | 174894 |
| J19/20 | VEGF | Vascular endothelial growth factor | - | 480 | 41250 | 37310 | 43174 | 40470 |
| | | | | | 41914 | 34458 | 34662 | 33866 |
| J21/22 | CCN4 | WNT1-inducible signaling pathway protein 1 | WISP-1 | 480 | 75382 | 76270 | 67422 | 85586 |
| | | | | | 80590 | 74150 | 65662 | 88930 |
| J23/24 | Negative control | Background | - | 480 | | | | |

**Supplementary Table 5.** Model summary for odds of 'large for gestational age' (LGA) fetal weight outcome

| | Model: LGA ~ diet*sex + (1ldam) | | | |
|---|---|---|---|---|
| **Fixed Effects** | *Odds Ratios* | *CI* | *t* | *p* |
| Intercept | 0.1 | 0.0 – 0.4 | -3.2 | **0.001** |
| HFHS Diet | 5.9 | 1.0 – 34.1 | 2.0 | **0.048** |
| Male Sex | 1.3 | 0.2 – 7.3 | 0.3 | 0.734 |
| Interaction | 0.3 | 0.0 – 2.4 | -1.1 | 0.259 |
| **Random Effects** | | | | |
| $\sigma^2$ | 3.29 | | | |
| $\tau_{00\ \mathrm{Dam}}$ | 1.17 | | | |
| ICC | 0.26 | | | |
| $N_{\mathrm{Dam}}$ | 20 | | | |
| Observations | 153 | | | |
| Marginal $R^2$ / Conditional $R^2$ | 0.101 / 0.337 | | | |

CI = 95% Confidence interval
*t* = t-statistic
*p* = p-value
$\sigma^2$ = sample variance
$\tau_{00\ \mathrm{Dam}}$ = between group (dam) variance
ICC = Interclass correlation coefficient
$N_{\mathrm{Dam}}$ = number of dams

**Supplementary Table 6.** Model summary for odds of 'small for gestational age' (SGA) fetal weight outcome

| | Model: SGA ~ diet*sex + (1ldam) | | | |
|---|---|---|---|---|
| **Fixed Effects** | *Odds Ratios* | *CI* | *t* | *p* |
| Intercept | 0.1 | 0.0 – 0.4 | -3.2 | **0.001** |
| HFHS Diet | 2.4 | 0.4 – 15.3 | 0.9 | 0.358 |
| Male Sex | 1.4 | 0.3 – 7.5 | 0.4 | 0.710 |
| Interaction | 0.4 | 0.1 – 3.5 | -0.8 | 0.420 |
| **Random Effects** | | | | |
| $\sigma^2$ | 3.29 | | | |
| $\tau_{00\ Dam}$ | 1.40 | | | |
| ICC | 0.30 | | | |
| $N_{Dam}$ | 20 | | | |
| Observations | 153 | | | |
| Marginal $R^2$ / Conditional $R^2$ | 0.023 / 0.314 | | | |

CI = 95% Confidence interval

*t* = t-statistic

*p* = p-value

$\sigma^2$ = sample variance

$\tau_{00\ Dam}$ = between group (dam) variance

ICC = Interclass correlation coefficient

$N_{Dam}$ = number of dams

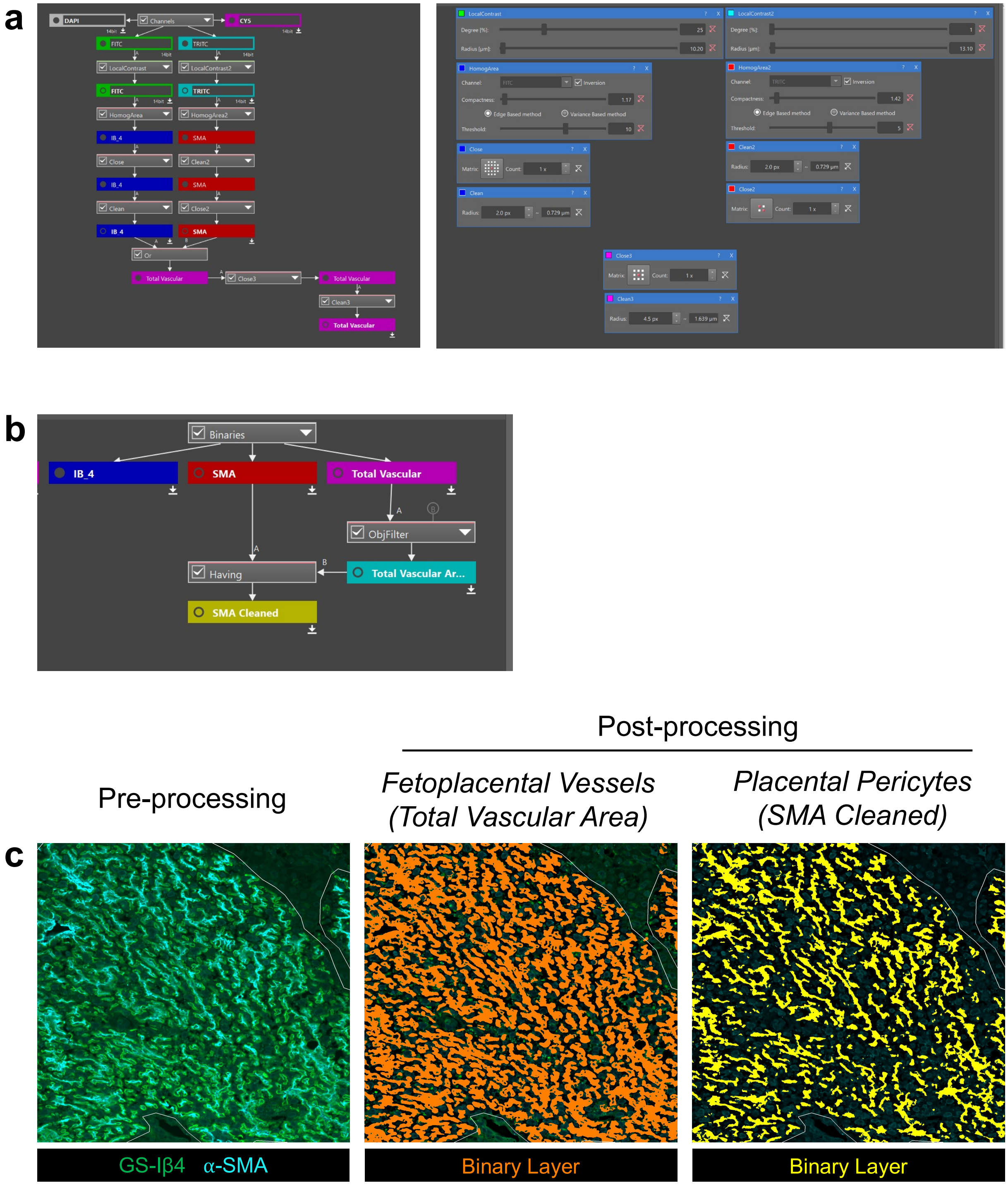

**Supplementary Figure 1. Automated image analysis pipeline for determination of placental vessel areas and microvascular stability. (a)** NIS Elements General Analysis 3 (GA3) image processing pipeline sequence for image pre-processing and binary layer generation (left), with respective parameters of each function (right). **(b)** Sequence of processing and cleaning for final development of binary layers for fetal vascular area (Total Vascular Area (TVA)) and placental pericytes (SMA cleaned), used in data extraction. **(c)** Representative images of image input (left panel, pre-processing) and binary layers of fetoplacental vessels (middle panel) and pericytes (right panel) generated for analysis following image processing. White outline represents region of interest (ROI) dividing placental labyrinth and junctional zones, only objects falling within the labyrinth ROI were included in analyses.

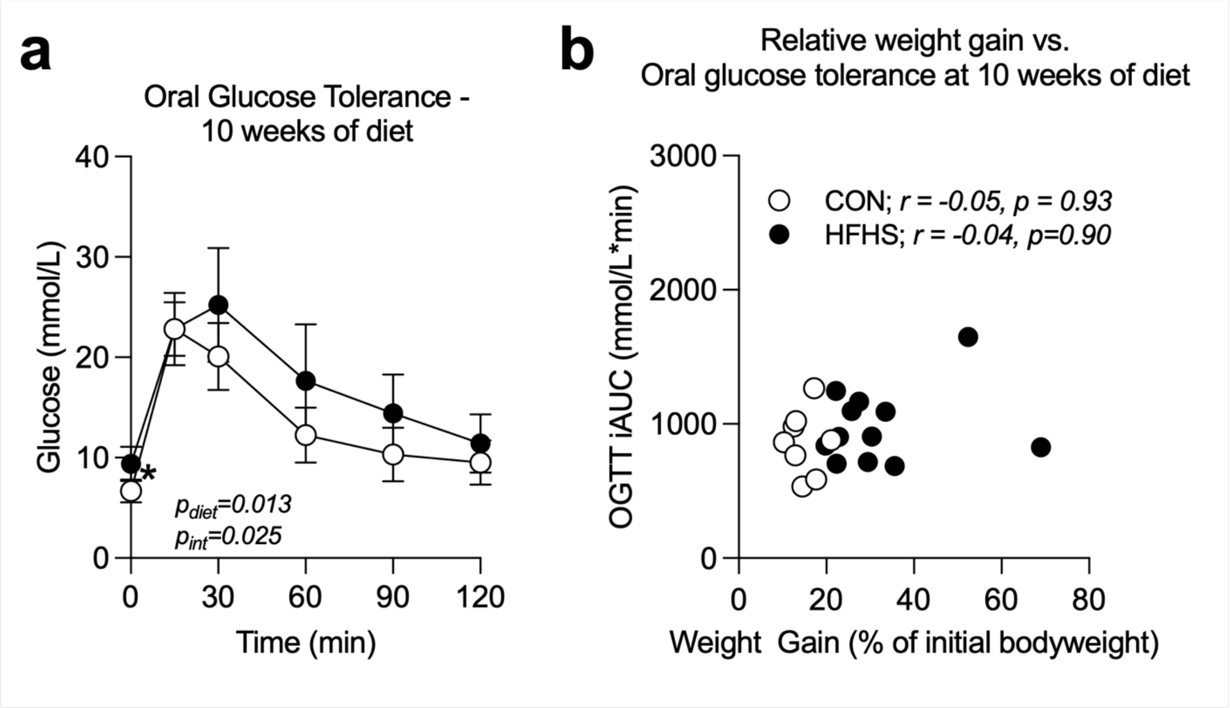

**Supplementary Figure 2. Absolute values of maternal blood glucose during OGTT and correlation of the incremental AUC with preconception weight gain at 10 weeks of dietary intervention. (a)** After 10 weeks, HFHS diet female mice were hyperglycemic before and throughout an oral glucose tolerance test, and showed an altered pattern of glucose clearance, but did not exhibit changes in their ability to clear a glucose bolus (incremental area under the curve) **(b)** Glucose clearance measured by the incremental area under the curves in (a) did not correlate with weight gained over the course 10-weeks of diet. as mean ± standard deviation (SD) in (a). Data points in (b) represent individual dams. Data were analyzed through repeated-measures two-way ANOVA, with posthoc Bonferroni-corrected pairwise comparisons (a) or Spearman-rank correlation (b). P-values from two-way ANOVA are indicated where main effects were statistically significant; Spearman's correlation coefficient (r) and associated p-values for each group are depicted in (b). * indicates $p<0.05$.

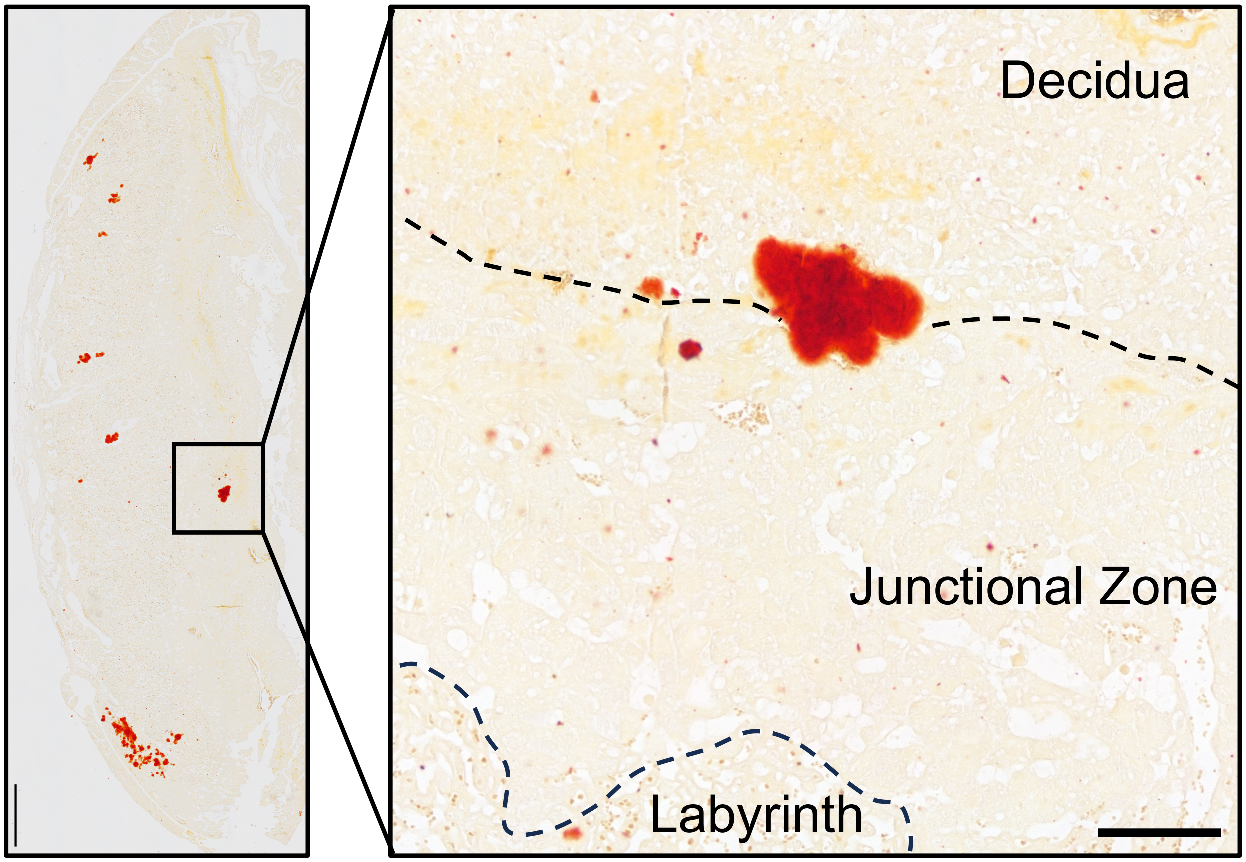

**Supplementary Figure 3. Occurrence of extra-labyrinthine calcification in HFHS placentas. (a)** Infrequent calcifications were observed in a small number of HFHS placental sections outside of the labyrinth. **(b)** Inset image from (a) shows a higher magnification of a calcified region at the intersection of the junctional zone and decidua, potentially affecting a spiral artery/arterial canal or outflowing venous channel. Scale = 500μm in (a) and 100 μm in (b).

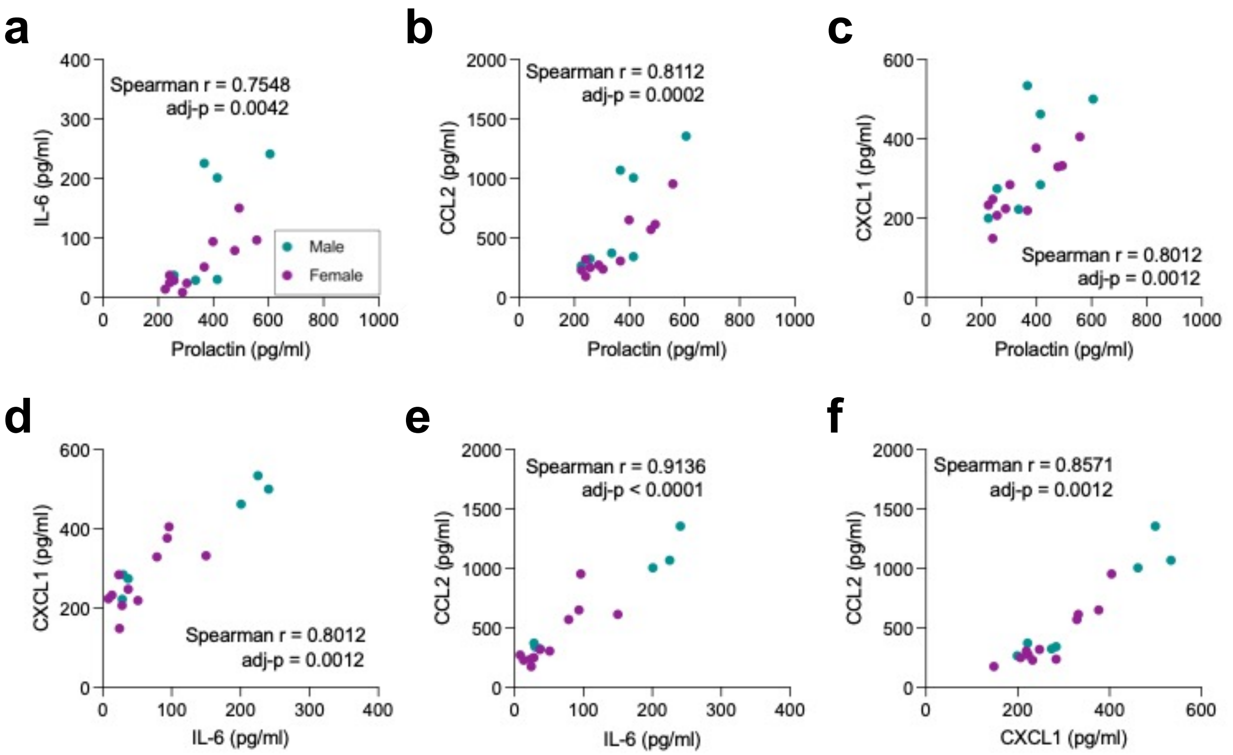

**Supplementary Figure 4. Circulating cytokine correlations in HFHS fetuses.** Serum cytokines found to be significantly elevated in HFHS female (magenta) and male (teal) fetuses were all significantly correlated with one another: **(a)** Prolactin vs. Il-6. **(b)** Prolactin vs. CCL2. **(c)** Prolactin vs. CXCL1. **(d)** IL-6 vs. CXCL1. **(e)** IL-6 vs. CCL2. **(f)** CXCL1 vs. CCL2. Data are presented as individual data points for each pooled fetal serum sample. Data were analysed using Spearman-rank correlation and p-values corrected for multiple testing using Bonferroni's method. Correlation coefficients (r) and adjusted p-values are depicted for each comparison.
